# Concurrent metacognitive evaluation is dissociable from perceptual evidence accumulation in the parietal cortex

**DOI:** 10.64898/2026.11.01.759033

**Authors:** Xiaodong Li, Yilin Chen, Ruixin Su, Tianming Yang

## Abstract

Adaptive behavior requires not only accumulating evidence toward a choice but also monitoring its reliability in real time. A central question is whether such concurrent uncertainty is read out directly from the circuit that accumulates perceptual evidence, or is computed by a separate process that runs alongside it. We trained two macaques on a perceptual discrimination task in which they could abandon deliberation at any moment and request assistance, so that uncertainty responses were in parallel with, rather than appended to, evidence accumulation. Behaviorally, a race between distinct perceptual and uncertainty accumulators outperformed models in which the opt-out decision was derived from the perceptual decision variable alone. To ask where the second accumulator resides, we recorded single units in the lateral intraparietal area (LIP), a canonical sensory-motor transformation area, and compared population dynamics with recurrent neural networks instantiating two architectures: one in which uncertainty is computed within the perceptual accumulator, and one in which it is computed outside and triggers the uncertainty response there. LIP exhibited all of these signatures for perceptual choices and none for uncertainty responses. Uncertainty-related activity rose late and stereotypically, lacking the coherence-scaled ramp of an accumulator, and separable from the perceptual accumulation. They are all predicted by the segregated architecture. Concurrent uncertainty is thus represented in LIP but is unlikely to be computed within LIP, indicating that metacognitive monitoring during deliberation is a dissociable operation rather than a derivative readout of the LIP perceptual circuit; its anatomical substrate is not localized here.

## Introduction

Adaptive behavior requires more than choosing among options. An agent must also evaluate whether the evidence in hand is sufficient to justify commitment. When that evaluation returns high uncertainty, the agent can seek more information, ask for help, or defer, rather than commit to a report it is likely to regret. This capacity to monitor uncertainty and act on it is a defining expression of metacognition^1,2^ and is observed across species, including macaque monkeys and humans^3–8^, where subjective confidence directly governs information seeking^9^.

How the brain implements this evaluation remains unresolved, and the debate can be sharpened into two logically separable questions. The first concerns computation: is the metacognitive signal a direct readout of the perceptual decision variable (DV), or is it the product of a dedicated decision process that accumulates evidence bearing on uncertainty in its own right? The readout view follows naturally from signal-detection and bounded-accumulation accounts, in which the state of the accumulator, together with elapsed time, is in principle sufficient to specify the probability of being correct; confidence is then a deterministic function of a quantity the perceptual system already computes ^4,10–12^. The dedicated-process view instead holds that uncertainty is evaluated by a distinct, partly independent computation. It is a second-order process operating on, but not identical to, the first-order evidence stream^13^. The second question follows and concerns the neural substrates: if a dedicated process exists, is it implemented within the same sensorimotor circuitry that forms the perceptual choice^4,12,14–19^, or in a separate circuit whose output is sent to and modulates the first-order decision-making circuitry^20–27^?

Resolving these questions is critical for our understanding of metacognition, yet progress has been limited by the paradigms available in animals for investigations at the level of identified neurons and circuits. Most rely on post-decision wagering or retrospective confidence reports^4,11,22,28,29^. Because the metacognitive report in these designs follows commitment to a choice, uncertainty-related activity is necessarily correlated with the terminal state of the perceptual accumulator, and the two accounts above become nearly indistinguishable. A genuine second-order computation and a passive readout predict similar retrospective correlations^30,31^. What is needed is a paradigm in which the metacognitive evaluation unfolds concurrently with, and can be committed to independently of, the perceptual decision.

Lateral intraparietal area (LIP) provides a well-motivated setting for such a test. LIP is among the best-characterized cortical substrates of bounded evidence accumulation. Single neurons in LIP show ramping activity whose slope scales with motion strength and whose dynamics predict both choice and reaction time ^32–35^. LIP activity also carries information relevant to metacognition. Firing rates at the end of motion viewing predict whether a monkey will subsequently select a post-decision sure-bet target^4^, and in a peri-decision wagering task, choice and wager signals are tightly co-registered across the LIP population during motion viewing, with near-zero temporal lag^19^. Such findings establish that the information required for a metacognitive judgment is present in LIP, and that it is available before commitment. They do not, however, establish how metacognitive judgments are computed and whether LIP performs the uncertainty computation. The ambiguity is particularly acute when the metacognitive report is conjunctively bound to a saccade target, as in wagering designs. Any uncertainty signal must then appear within LIP’s spatial-motor representation regardless of where it was generated.

We therefore adopted a concurrent uncertainty-estimation task^36^. Monkeys judged the direction of random-dot motion and could, at any moment before committing to a direction, request assistance by choosing an opt-out option. Because the uncertainty response is available throughout deliberation and is voluntarily timed, it yields its own reaction-time distribution and coherence dependence as the behavioral fingerprints of its decision process. In addition, because it is spatially decoupled from the perceptual saccade, we may examine the uncertainty-related evidence accumulation process in LIP that cannot be attributed to that for the perceptual choice.

We find that LIP displays the canonical hallmarks of a bounded accumulator for the perceptual decision, but not for the uncertainty response. Both the neurons’ firing rates and their trial-to-trial variability structure are inconsistent with a locally implemented accumulation-to-bound process. Population-geometry analyses and second-order statistics instead indicate that the uncertainty signal is injected into LIP from an external source. These results argue that first-order perceptual decisions and their concurrent metacognitive evaluation depend on dissociable computations implemented in distinct circuits, and they challenge the theory that metacognitive signals are a direct readout of the decision variable in LIP.

## Results

### A concurrent uncertainty-estimation task

Two monkeys were trained to perform a concurrent uncertainty estimation (CUE) task (**Fig. 1a**). On each trial, monkeys discriminated the net direction of a random-dot motion stimulus by saccading to one of two perceptual decision targets, in a reaction-time version of the classic motion discrimination task^33^. Motion direction was aligned to one of the two targets, and coherence was drawn randomly from 0%, 6.4%, 12.8%, 25.6%. Crucially, an uncertainty response (UR) target was available throughout the stimulus epoch. Selecting the UR target triggered the presentation of a second stimulus moving in the same direction as the first but at a high coherence (25.6%), after which the monkey could report its direction with ease. Correct direction reports yielded the same reward whether the UR target had been selected, whereas errors incurred a timeout punishment. Because the UR option carried an additional time cost but no reward bonus, it was advantageous only when the uncertainty about the initial stimulus was high. As a control, monkeys were also tested on a standard random-dot motion direction discrimination task (STD) in which the UR target was unavailable.

**Fig. 1.**
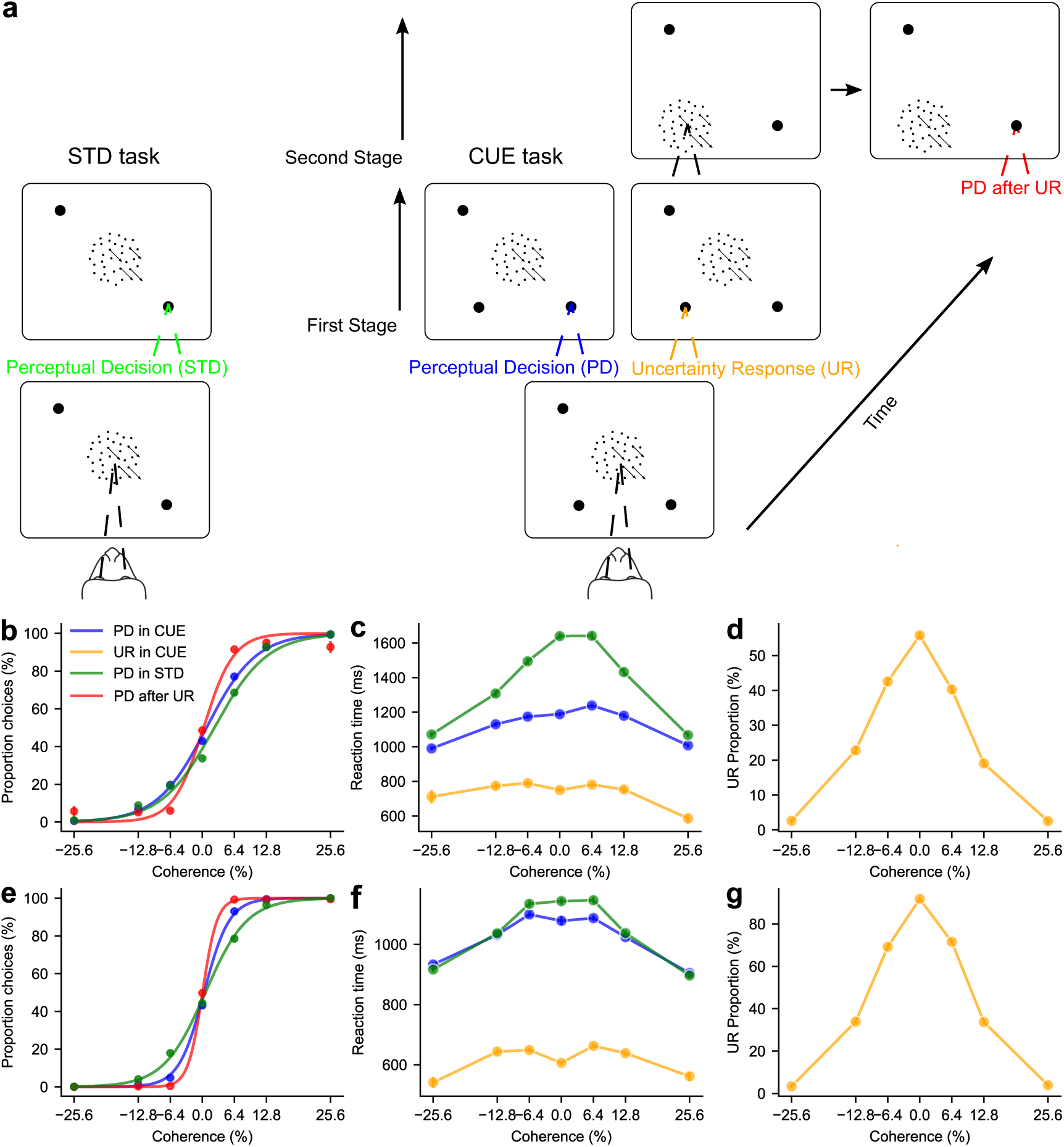
Behavior paradigm and performance. **a.** Schematic of the concurrent uncertainty estimation (CUE) task. Each trial began with the monkey fixating on a central point (top). After a 500-ms delay, two perceptual decision (PD) targets (left/right, indicating motion direction) and an uncertainty response (UR) target appeared, followed by a variable delay and the onset of a random-dot motion stimulus. The monkey could either (i) saccade directly to one of the two PD targets to report motion direction (first stage, left branch), ending the trial with reward for a correct report or a timeout punishment for an error; or (ii) saccade to the UR target (right branch), which extinguished the first stimulus and triggered a second stage in which a new random-dot stimulus moved in the same direction at the highest of the four (25.6%). Afterwards, the monkey reported the direction. Correct direction reports were rewarded equally regardless of whether the UR target had been selected. The standard (STD) control task was identical except that the UR target was not available. **b-d.** Behavioral performance of Monkey F (36 units; CUE task 27277 trials; STD task 13588 trials), plotted as a function of first-stage random-dot signed coherence (+: right; -: left). **b.** Psychometric curve for perceptual decisions. Green, PD in the STD task (PD in STD); blue, PD in the CUE task on trials in which the UR target was not chosen (PD in CUE); red, PD in the second stage of the CUE task, after the UR target had been chosen. **c.** Reaction times for the first-stage decisions. **d.** Proportion of trials on which the monkey selected the UR target. **e-g.** Same as **b-d** for Monkey M (55 units; CUE task 43877 trials; STD task 10086 trials).

Two features of this design are essential to the questions we set out to answer. First, the uncertainty response is available throughout deliberation and is voluntarily timed, so it generates its own reaction-time distribution and its own coherence dependence. These behavioural fingerprints cannot be provided by retrospective wagering designs. Second, the UR target is spatially dissociated from the two perceptual targets, so uncertainty-related activity at the corresponding location cannot be attributed to the developing perceptual choice.

Both monkeys’ first-stage perceptual choices showed the canonical dependence on motion strength: accuracy increased monotonically with coherence in both the CUE and STD tasks (**Fig. 1b, e**, blue and green). At matched coherence, first-stage perceptual decisions in the CUE task were more accurate than those in the STD task (Mann-Whitney U one-sided test *p* < 1 × 10^−1^^6^, Monkey F; *p* < 1 × 10^−1^^6^, Monkey M), consistent with selective opt-out on the highly uncertain trials. When monkeys selected the UR target and viewed the second-stage high-coherence stimulus, their subsequent direction reports were near-ceiling and independent of the first-stage coherence (**Fig. 1b, e**, red), confirming that the UR option provided effective assistance.

The probability of choosing the UR target tracked stimulus difficulty. It was highest at 0% coherence, declined monotonically with increasing coherence, and approached zero at 25.6%, where the second stimulus would offer no informational advantage (**Fig. 1d, g**). Thus, the monkeys took advantage of the UR option strategically rather than indiscriminately.

### The uncertainty response is emitted concurrently with the perceptual decision

Reaction times revealed a striking dissociation between the two response types. First-stage perceptual decisions exhibited the classical inverse relationship with coherence in both tasks (**Fig. 1c, f** blue and green). In addition, the reaction times showed a dip at 0 coherence in the CUE task. UR latencies, in contrast, did not vary monotonically with coherence. They were shorter at the two extremes (0 and 25.6%) and longer at intermediate coherences (**Fig. 1c, f**, orange; Mann-Whitney U one-sided test *p* = 8.194 × 10^−9^, Monkey F; *p* = 3.440 × 10^−5^^2^, Monkey M).

Three features show that the UR is not a readout appended after a completed perceptual decision. First, uncertainty responses were faster than first-stage perceptual decisions at every coherence (Linear Mixed model, *H*_0_ = *UR* ≥ *PD*, Monkey F: *β* = 429.4, *se* = 3.9, *z* = 109.66, *sided p* < 1 × 10^−1^^6^; Monkey M: *β* = 410.8, *se* = 2.4, *z* = 172.03, *sided p* < 1 × 10^−1^^6^). In a purely sequential scheme, the UR could not finish before the perceptual decision it follows, so this rules out pure sequential processing. Second, the non-monotonic coherence dependence of UR latency is incompatible with any scheme in which the uncertainty response inherits its timing from a completed perceptual choice, since perceptual latencies decrease monotonically with coherence (**Fig. 1c, f**). Lastly, first-stage perceptual decisions in the CUE task were faster than those in the STD task (Linear Mixed model, *H*_0_ = *PD* ≥ *STD*, Monkey F: *β* = 197.6, *se* = 4.6, *z* = 43.34, *sided p* < 1 × 10^−1^^6^; Monkey M: *β* = 15.9, *se* = 2.9, *z* = 5.47, *sided p* = 2.296 × 10^−8^), arguing against a serial scheme in which the monkey first evaluates whether to opt out and then, conditional on staying in, commits to a perceptual decision.

Taken together, these features rule out pure serial processing. The UR is emitted concurrently with, not downstream of, a completed perceptual decision. This is a statement about timing, not about computation. A fast, concurrent UR is equally compatible with a concurrent readout of the perceptual accumulator, in which an inner bound is crossed early, and with a dedicated second accumulator that races the perceptual process in parallel. Latency data alone cannot even exclude a short shared perceptual phase followed by an independent UR choice. Whether the UR is computed by a dedicated second-order process, or is instead read out from the perceptual accumulator, therefore remains open, and latencies alone cannot resolve it. So we fit the two competing model classes to the joint structure of accuracy, latency, and opt-out rate, and ask which better accounts for these data (**Fig. 2**).

**Fig. 2.**
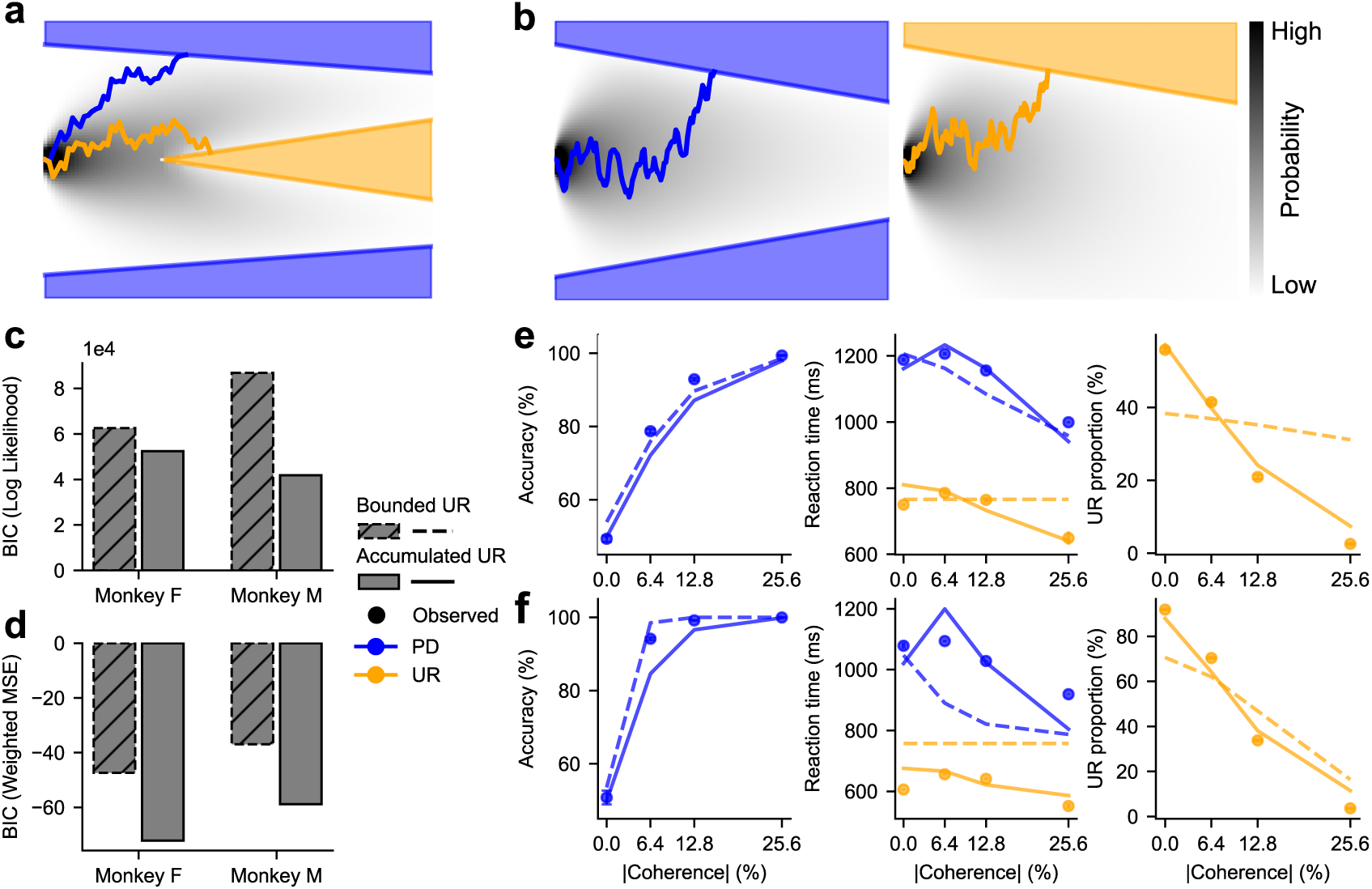
A parallel accumulator for the uncertainty response accounts for behavior better than an inner decision bound. **a, b.** Schematics of the two competing models. In both, a single accumulator drives the perceptual decision. The models differ in how the UR is generated. In the single-accumulator model (**a**), the UR (orange) is read out from the perceptual accumulator itself, as an inner bound is crossed. This single-accumulator model is a simplification of the optimal policy model in the previous study^36^. In the parallel-accumulator model (**b**), the UR is generated by a separate accumulator that evolves concurrently with the perceptual accumulator and whose drift rate decreases with coherence (a segregated scheme). A UR is emitted when this independent accumulator reaches its bound first. **c.** Bayesian information criterion (BIC) for the two models, fit to the pointwise likelihood of the reaction-time distributions, for Monkey F (36 units, 27277 trials) and Monkey M (55 units, 43877 trials). Lower is better. **d.** Same as c, but for the weighted mean squared error (wMSE) computed jointly over accuracy, reaction time, and UR proportion. By both criteria and in both monkeys, the parallel-accumulator model provides the better account. **e, f.** Model fits (wMSE method) for accuracy (left), reaction time (middle), and UR proportion (right), for Monkey F (**e**) and Monkey M (**f**), respectively. Points, data; dashed lines, single-accumulator model; solid lines, parallel-accumulator model. Blue, PD option; orange, UR option.

### A dedicated uncertainty response accumulator accounts for behavior better than a readout of the perceptual decision variable

To adjudicate between the readout and dedicated-process accounts, we fit two drift-diffusion models that differ only in the origin of the signal supporting the uncertainty response (**Fig. 2a, b**). Both models describe the perceptual decision with a single accumulator whose drift rate scales with signed coherence, and both include a shared non-decision time. They diverge in how the UR is produced.

In the single-accumulator model (**Fig. 2a**), the uncertainty response is generated within the perceptual accumulator. A UR is emitted when the accumulated evidence lingers near zero and crosses an inner bound. Thus, a single evidence stream jointly determines both option choice and the decision to opt out. This model follows the optimal-policy formulation^36^ in treating the uncertainty as an inner region of the perceptual decision variable and instantiates the readout hypothesis.

In the parallel-accumulator model (**Fig. 2b**), the uncertainty response is generated by a distinct accumulator that races against the perceptual accumulator. Its drift rate decreases with coherence, so that weak or absent motion drives it toward its bound while strong motion drives it away. This instantiates the dedicated-process hypothesis, in which the signal supporting the uncertainty response is computed by a process distinct from the perceptual decision, although both are fed by the same sensory evidence.

We fit each model to the behavior of both monkeys, using two complementary objectives: the pointwise likelihood of the full reaction-time distributions (**Fig. 2c**) and a weighted mean squared error (wMSE) computed jointly over accuracy, reaction time, and UR proportion (**Fig. 2d**). The parallel-accumulator model was favored under both criteria in both monkeys: BIC on the pointwise reaction-time likelihood (**Fig. 2c**; Monkey F: 61346.38 vs 2442.93, *N* = 27277 ; Monkey M, 86804.23 vs 41801.30, *N* = 43877) and BIC on the wMSE (**Fig. 2d**, Monkey F: - 51.89 vs -72.2, *N* = 16; Monkey M, -36.97 vs -58.81, *N* = 16, **methods**).

Critically, the two models made divergent predictions about the signature that first motivated them. The single-accumulator model, in which UR timing is tied to the perceptual accumulator, could not reproduce the non-monotonic dependence of UR and PD latency on coherence, predicting a monotonic profile instead. In particular, the parallel-accumulator model captured the slightly faster RT of PD at zero coherence than at the coherence of 12.8%. The UR accumulator’s drift is highest at zero coherence, producing fast URs, which truncate the slow tail of the PD distribution (**Fig. 2e, f**, middle). The parallel-accumulator model also reproduced the elevated accuracy of first-stage perceptual choices and the monotonic decline of UR proportion with coherence, whereas the single-accumulator model systematically misfits one or more of these measures (**Fig. 2e, f**).

Thus, at the algorithmic level, behavior is better explained when the uncertainty response arises from a dedicated accumulator operating in parallel with the perceptual decision than when it is read out from the perceptual accumulator itself. This answers the first of the two questions posed in the introduction and localizes the second rather than answering it. A dedicated accumulator must be implemented somewhere. Two possibilities remain. If the implementation is local, the uncertainty accumulator is housed within the same sensory-motor transformation circuitry that forms the perceptual choice. Alternatively, the uncertainty accumulator may reside outside this circuitry, and we should observe an absence of uncertainty signal within the sensory-motor transformation circuitry.

Area LIP is the natural place to begin, both because its ramping activity is the canonical correlate of perceptual evidence accumulation^32–35^ and because firing rates there already predict metacognitive reports^4,19^. We recorded from LIP to distinguish these possibilities.

### LIP accumulates evidence for perceptual decisions but not for uncertainty responses

We recorded single-unit activity from LIP in both monkeys while STD and CUE tasks were interleaved in alternating blocks. For each unit, we mapped the response field with a memory-guided saccade task. Of the 99 (Monkey F) and 118 (Monkey M) LIP units recorded, 36 (F) and 55 (M) units showed significant spatial selectivity and were used in the behavioral and model-fit panels (**Fig. 1 and 2**); 30 (F) and 49 (M) units additionally exceeded 600 trials and were retained for all subsequent analyses.

In the STD task, we placed one of the perceptual choice target inside the response field of the recorded neuron (**Fig. 3a,d**). On CUE trials, we analyzed each neuron under two target configurations: with a perceptual target placed inside its response field (PD configuration; **Fig. 3b,e**) and, on separate trials, with the UR target inside its response field (UR configuration; **Fig. 3c,f**).

**Fig. 3.**
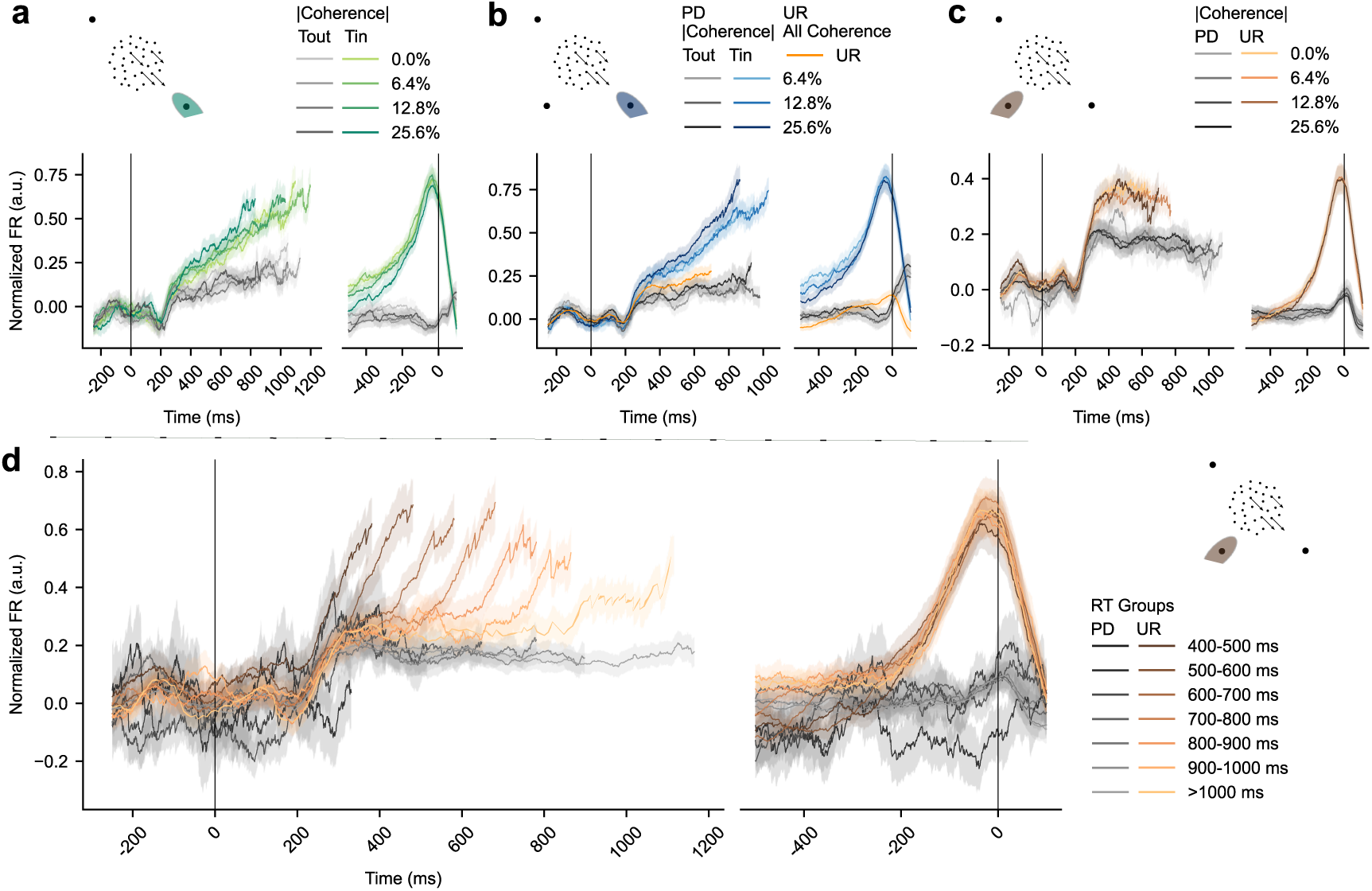
LIP responses across receptive-field configurations and tasks. Peristimulus time histograms (PSTHs) of LIP activity, pooled across recorded units from both monkeys. Each unit was normalized before averaging, and units were weighted equally. Within each panel, responses are aligned to motion onset (left) and to saccade onset (right). **a**. The STD task. PSTHs grouped by coherence (color intensity indicates coherence; see key), pooled across both monkeys. **b**. The CUE task with one of the perceptual-decision targets in the response field. PSTHs grouped by coherence. **c**. The CUE task with the UR target in the response field. PSTHs grouped by coherence. **d**. The CUE task with the UR target in the response field. PSTHs grouped by reaction time. Conditions with too few trials (**b**, 0.0% in PD, Monkey M: *T*_*in*_95 trials in 42 units, *T*_*out*_108 trials in 38 units; **c**, 25.6% in UR, Monkey F: 62 trials in 15 units; Monkey M: 120 trials in 36 units; **d**, <400ms in UR, Monkey F: 56 trials in 8 units; Monkey M: 268 trials in 42 units) are not plotted here. For the motion-onset-aligned traces, each condition was truncated at the across-unit average of the single-unit median RTs. The UR condition in **b** was truncated at three-quarters of that value, because UR reaction times were substantially shorter than PD reaction times. Colored lines except orange in **b**, *T*_*in*_trials; Grey lines, *T*_*out*_trials. Individual monkeys showed similar patterns (**Extended Data Fig. 1 & 2**).

In the STD task, LIP activity exhibited the canonical ramping pattern. Neurons’ firing rates diverged shortly after motion onset, ramping upward when motion favored the target inside the response field (T_in_) and downward when it favored the opposite target (T_out_), faster with higher coherence (**Fig. 3a**; individual monkeys: **Extended Data Fig. 1a & 2a**). A similar pattern was observed on PD-choice trials in the CUE task, in which the monkey selected a perceptual target directly (**Fig. 3b**; individual monkeys: **Extended Data Fig. 1b & 2b**), confirming that the recorded neurons were a representative sample of evidence-accumulating LIP cells and that the presence of the UR option did not abolish the perceptual accumulation signal.

On trials in the PD configuration in which the monkey selected the UR target, LIP activity followed an intermediate trajectory between the T_in_ and T_out_ perceptual-choice traces (**Fig. 3b**, orange). This is the classical neural signature of low-confidence trials^4^: when accumulated evidence is ambiguous, neither perceptual bound is reached, and the subject opts out. Viewed from the perceptual accumulator, the UR is selected precisely when perceptual evidence is weakest.

We next turned to the UR configuration to ask the converse, and more decisive, question: do these same neurons accumulate evidence *for* the uncertainty response when the UR target falls in their response field? If the UR accumulator is hosted locally in LIP, we expected the textbook signatures of accumulation. There should be ramping toward saccade onset when UR, T_in_ in this configuration, is chosen; the ramping activity strength should be modulated by coherence, with steeper ramping when evidence for UR is stronger, that is, at lower coherences; and we should see ramping downward from baseline on trials in which the monkey rejected the UR target in favor of a perceptual choice (T_out_).

None of these signatures were present (**Fig. 3c**; individual monkeys: **Extended Data Fig. 1c & 2 c**). Grouped by coherence, UR-choice trials showed an elevated mean firing rate relative to perceptual-choice trials, which is expected as the UR target was in the receptive field. However, there was no systematic ramping during the stimulus epoch and no separation across these coherence levels. On trials in which the monkey rejected the UR and made a first-stage perceptual choice, firing rates remained near baseline rather than ramping downward, as they should if these neurons encoded a UR decision variable. Moreover, when incorporating the rare trials in which monkeys chose UR at 25.6% coherence (**Extended Data Fig. 6**), the recorded units showed substantially higher firing rates compared with those observed at other coherence levels. These findings are inconsistent with the prior hypothesis.

One concern with grouping by coherence is that UR reaction times were quite variable and relatively flat across coherences (**Fig. 1d, g**), which could mask a motion-strength-dependent ramping. We therefore re-grouped trials by reaction time, reasoning that a fast accumulator should yield steeper ramps on short-RT trials. This regrouping revealed a qualitatively distinct pattern (**Fig. 3d**; individual monkeys: **Extended Data Fig. 1d & 2d**). On UR-choice trials, firing rates remained near baseline regardless of RT and then rose abruptly in the final ∼300 ms before the saccade, in a stereotyped, RT-independent manner. This presaccadic rise is consistent with a saccade-related burst rather than evidence accumulation, because a genuine accumulation signal produces more gradual ramps whose slope varies with RT, rather than a stereotyped rise locked to the saccade. Applying the same RT-grouping to the STD task or to the PD-configuration trials in the CUE task reproduced the classical RT-dependent ramping (**Extended Data Fig. 3**), confirming that the absence of ramping in the UR configuration is not an artifact of the RT grouping.

Together, these results show that although LIP neurons carry the canonical accumulation signal for perceptual choices, including the classical low-confidence signature on opt-out trials, they do not accumulate evidence for the uncertainty response itself. The dedicated UR accumulator favored by our behavioral model thus appears to be dissociable from LIP’s perceptual accumulation circuitry, strongly suggesting that its neural substrate resides elsewhere.

### Two recurrent network agents instantiate local and remote implementations

Interpreting the dissociation above requires a benchmark. The abrupt, RT-independent presaccadic rise on UR trials is consistent with a signal arriving from outside LIP, but it is not diagnostic on its own. We therefore need to know what each architecture looks like when measured the way we measured LIP. We instantiated both as recurrent neural network (RNN) agents trained with reinforcement learning to perform the CUE task, applied identical analyses to their unit activity, and asked which better reproduces the LIP findings.

Two agents are created. The integrated-UR (IUR) agent computes the UR with the same recurrent circuit that supports the perceptual choice. Note that a single decision module of this kind can implement either a single accumulator with an inner bound or a parallel within-circuit accumulator, so the IUR agent serves as the joint instantiation of both local architectures. The segregated-UR (SUR) agent instantiates the remote implementation model, in which the UR signal is computed outside the perceptual choice circuit, and the late rise in LIP-like units reflects a signal injected from an upstream module once that module has decided to opt out.

The two agents share the same architecture: a 48-unit vanilla RNN receiving four input streams: a fixation requirement, two channels of stimulus motion (left vs. right), and a UR cue (**Fig. 4a**), except that the SUR agent additionally receives an external UR trigger signal (**Fig. 4d**). The RNN produces an action via a softmax policy over a learned readout. Each trial comprises the inter-trial interval, fixation period, and first- and second-stage stimulus epochs, with the durations partly determined by the agent’s own actions. Agents were trained with proximal policy optimization using reward shaping that incentivizes accuracy for both first- and second-stage PD choices and penalizes fixation breaks, response failures, UR choices, and, in the SUR agent, premature URs. The training was curriculum-staged (**Methods**).

**Fig. 4.**
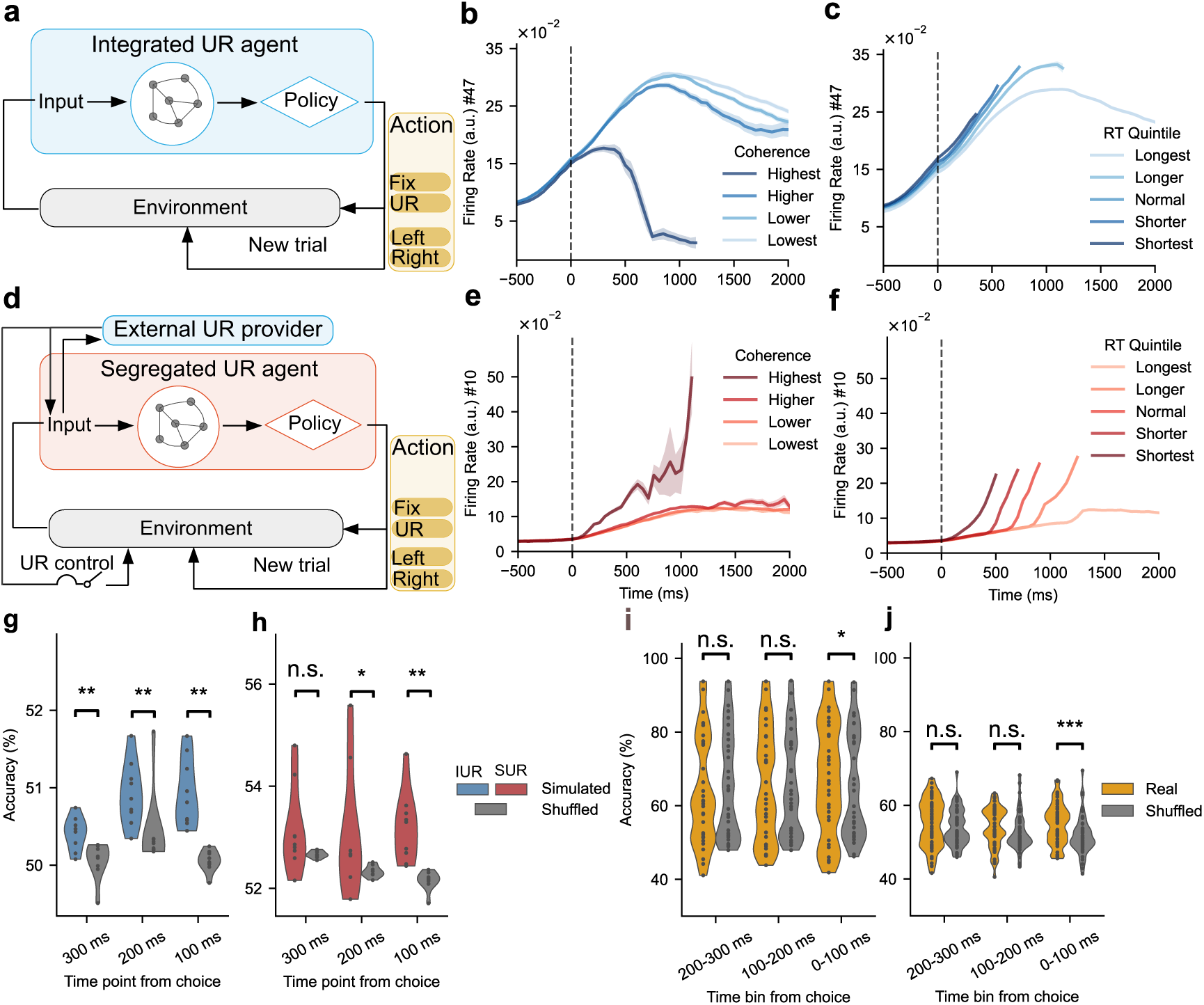
A segregated, but not integrated, RNN agent reproduces LIP unit activity and decoding profiles during UR choices. **a**. Schematic of the integrated UR agent (IUR). A 48-unit vanilla RNN receives four inputs from the environment at every time step: a fixation cue, stimulus motion (two channels), and a UR cue. It produces an action through a softmax policy. The agent is trained with proximal policy optimization, combining a policy loss, a value loss, and an entropy-regularization term that encourages exploration (**Methods**). **b**, **c**. Example UR unit in the trained IUR agent, with trials grouped by motion coherence (**b**) and by RT quintile (**c**). **d**. Schematic of the segregated UR agent (SUR). The decision RNN is identical to that in (a), except that it additionally receives a UR-trigger input from an external UR provider, which is a frozen, previously trained IUR agent receiving the same environmental inputs. The SUR agent is trained to emit a UR action only within 200ms of an external trigger, and UR actions outside this window are penalized (**Methods**). **e**, **f**. Example unit of the trained SUR agent. The units show a relatively flat baseline across RTs (**f**), followed by a stereotyped late rise, matching the LIP profile. Error bars are depicted with sem. **g**, **h**. kNN decoding of UR versus PD choices from positive UR units of the IUR agent (**g**, 9 units) and the SUR agent (**h**, 9 units). Colored distributions show the decoding accuracy; gray distributions show shuffled-label controls. P values are from one-sided Wilcoxon signed-rank tests (real > shuffled). **i**, **j**. The same decoding analysis applied to LIP UR-target units in monkey F (**i**, 30 units) and monkey M (**j**, 49 units). n.s. *p* > 0.05; * *p* < 0.05; ** *p* < 0.01; *** *p* < 0.001.

The agents differ in how the UR action is produced. The IUR agent has a single decision module that produces both PD and UR actions from the same RNN architecture. While for SUR agent, it is trained to emit a UR action only within a 200 ms window following a signal provided by an external UR provider, and UR actions outside this window are penalized. To match behavior between the two agents and isolate the architectural difference, we used the IUR agent as this external provider, frozen during the training of the SUR agent (**Methods**). This design precludes the SUR agent from generating UR internally (**Extended Data Fig. 4f**, No Ext.).

Both agents learned the task and reproduced the monkeys’ major behavioral patterns, including coherence-dependent UR choice proportions and faster UR reaction times compared with PD trials (**Extended Data Fig. S4b, e**). As enforced by training, the SUR agent emitted UR actions only following an external trigger (**Extended Data Fig. 4f** UR vs No Ext.).

One consequence of this design must be made explicit before the comparisons that follow. By construction, the SUR agent’s UR actions are confined to a window after the external trigger. The late timing of UR-related activity in the SUR network is therefore imposed rather than discovered, and the agreement between SUR single-unit profiles and LIP (**Fig. 4e, f**) should be read as a consistency check rather than as independent evidence. Analyses can be informative only when they are not constrained by this training signal. The following analyses are created with this point in mind.

### Only the segregated agent reproduces the LIP activity profiles

To compare single-unit activity across agents and LIP, we identified UR units in each agent as those whose activity makes a consistent positive contribution to the UR output, based on the Jacobian of the policy with respect to unit activations (**Methods, Extended Data Fig. 11**). These are the model analogues of LIP neurons whose response field covers the UR target.

Stratifying trials by motion coherence and by reaction time produced a clean double dissociation between the agents. IUR units are heterogeneous in firing patterns (**Fig. 4b**, **Extended Data Fig. 5a vs 5b** third column). When grouped by coherence, IUR units exhibited coherence-dependent ramping (**Fig. 4b**, **Extended Data Fig. 5a** third column), whereas SUR units showed a relatively flat, coherence-independent baseline, except for the highest coherence level that has an early rise (**Fig. 4e**, **Extended Data Fig. 5b** third column). A similar pattern is also observed in the LIP data (**Extended Data Fig. 6**). When grouped by reaction time, IUR units displayed RT-dependent ramping (**Fig. 4c**, **Extended Data Fig. 5a,** fourth column), whereas SUR units exhibited a stereotyped late rise atop an RT-independent baseline (**Fig. 4f, Extended Data Fig. 5b,** fourth column). In both cases, the SUR pattern is consistent with LIP activity during UR selection within the UR receptive field (**Fig. 3c, d**), whereas the IUR pattern is not.

Activity-level comparisons may understate the difference between models, because population-level information about the upcoming choice can be present even when individual units do not ramp. We therefore decoded UR versus PD choices from UR-unit population activity using a k-nearest-neighbors classifier (k = 5, **Methods**).

The decoding profiles recapitulate the activity-level dissociation (**Fig. 4g, h**). In the IUR agent, the UR choice was decodable significantly above chance throughout the stimulus epoch, indicating that the UR decision forms gradually within the network (**Fig. 4g**, Wilcoxon signed-rank test *H*_0_: Simulated>Shuffled. *sided p* = ( 5.859 × 10^−3^, 5.859 × 10^−3^, 1.953 × 10^−3^) from early to late). In the SUR agent, decoding accuracy was near chance early on and rose only in the last two time points before the saccade, indicating that the UR decision is not computed initially within the network, and the UR signal enters the decision RNN only late, through the external trigger (**Fig. 4h**, Wilcoxon signed-rank test *H*_0_: Simulated>Shuffled. *sided p* = ( 8.203 × 10^−2^, 4.883 × 10^−2^, 1.953 × 10^−3^)). A similar late emergence appeared in both monkeys’ LIP data (**Fig. 4i, j**; Wilcoxon signed-rank test *H*_0_: Real>Shuffled. Monkey F: *sided p* = (8.188 × 10^−1^, 3.688 × 10^−1^, 4.812 × 10^−2^); Monkey M: *sided p* = (1.007 × 10^−1^, 9.812 × 10^−2^, 3.592 × 10^−4^)), with the final pre-saccadic window alone.

### Variance and autocorrelation distinguish accumulation from late injection

In addition to the activity decoding analyses, the architectural difference between IUR and SUR also makes specific predictions about the second-order statistics of the neuronal responses that go beyond trial-averaged firing rates.

We tested these predictions using the variance of the conditional expectation (VarCE) and the correlation of the conditional expectation (CorCE)^37^ (**Methods**). VarCE estimates the variance of the latent conditional mean firing rate after subtracting the point-process variance expected from a doubly stochastic Poisson process, and CorCE estimates the corresponding autocorrelation between time bins. Each provides informative diagnoses on the computation in the network.

The predictions are opposite in sign. A unit whose activity reflects a within-network accumulator produces a VarCE that grows approximately linearly with time, because variance accumulates with each independent evidence increment. Correspondingly, its CorCE is high between nearby bins and decays gradually with lag, because the latent state at any time is the running sum of all earlier increments. In contrast, a unit whose activity reflects a late, externally injected signal produces a VarCE that is small early and rises rapidly once the external signal arrives, and a CorCE in which early bins decorrelate from late bins, because the late state is dominated by a signal largely independent of the early within-network state.

This is what we observed. In the IUR agent, VarCE for both UR and PD units rose gradually at comparable rates throughout the stimulus epoch (**Fig. 5a**), consistent with both populations performing within-network evidence accumulation. In the SUR agent, by contrast, VarCE for UR units rises more steeply and significantly exceeds that of PD-unit (**Fig. 5b**; one-sided Wilcoxon signed-rank test across units). LIP units show the same pattern: VarCE rises more rapidly when a UR target falls within their response field than when a PD target does (**Fig. 5c, d**), mirroring the SUR agent but not the IUR agent.

**Fig. 5.**
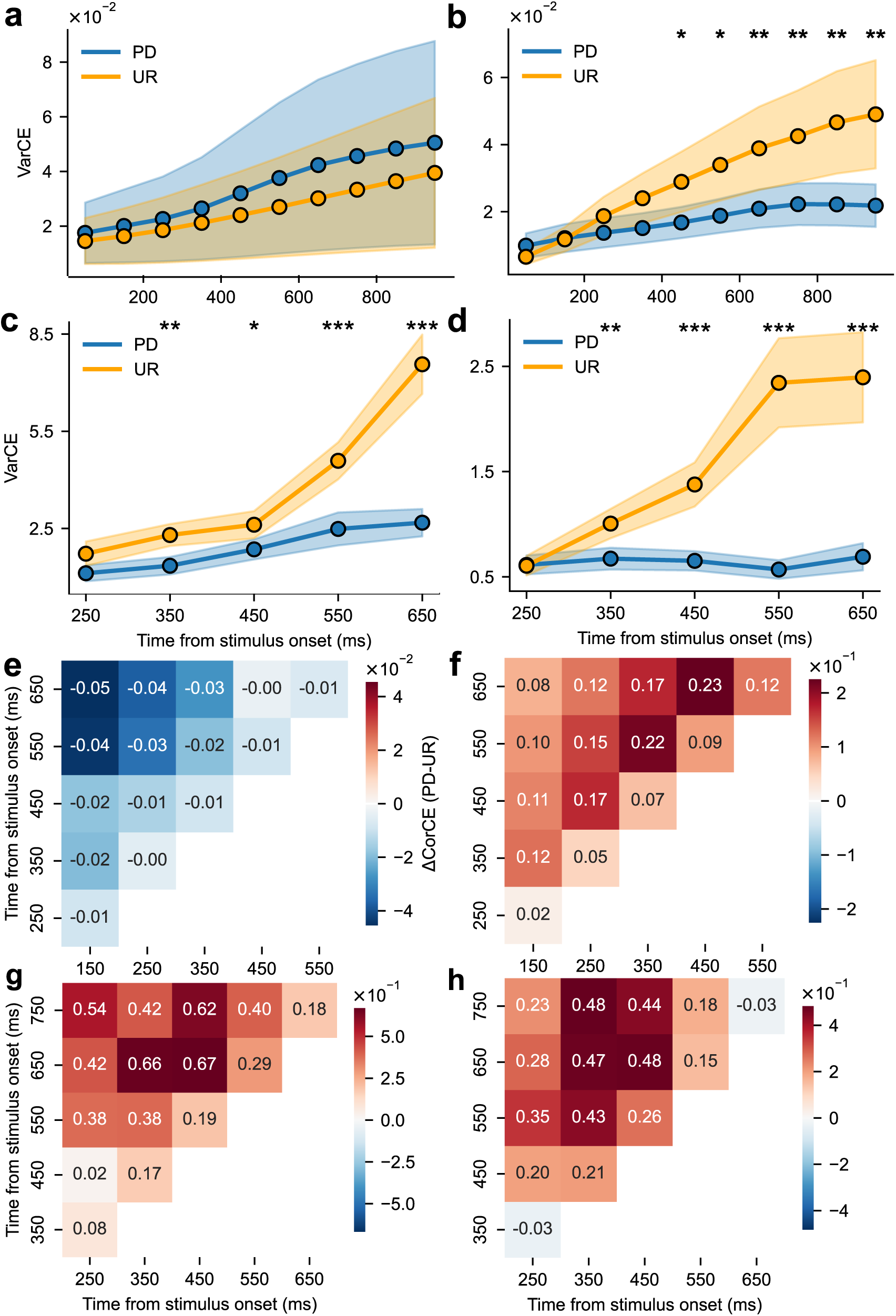
Variance and autocorrelation of UR-unit activity during the stimulus epoch. **a-d.** VarCE computed in successive time bins during the stimulus epoch for PD trials (blue) and UR trials (yellow), in positive UR units. **a.** IUR agent (9 units, 8 sessions), **b.** the SUR agent (9 units, 8 sessions), **c.** LIP units from monkey F (30 units), and **d.** LIP units from monkey M (49 units). Lines and shading show mean ± s.e.m. across the unit set. \**p*< 0.05, one-sided Wilcoxon signed-rank test. **e-h.** Difference in CorCE between PD and UR trials, computed between pairs of time bins in the same units. Red, stronger temporal autocorrelation on PD trials; blue, the reverse. Only the lower triangle is shown because CorCE is symmetric; the diagonal is omitted because CorCE on the diagonal is one by definition. Time advances from lower left to upper right.

We next computed CorCE separately for PD and UR trials and examined their difference (ΔCorCE = CorCE on PD trials minus CorCE on UR trials) at each pair of time bins (**Fig. 5e-h**; red, stronger autocorrelation on PD trials; blue, stronger on UR trials). In the IUR agent, this difference was small and unstructured (**Fig. 5e**), as expected when PD and UR trials are produced by the same within-network accumulator and share a temporal autocorrelation structure. In the SUR agent, the difference was large, especially in the off-diagonal entries linking early to late stimulus bins (**Fig. 5f**), reflecting that on UR trials the late state is dominated by the external trigger and is uncorrelated with the early within-network state, whereas on PD trials early and late bins remain coupled through the accumulator. LIP neurons in both monkeys showed the SUR pattern (**Fig. 5g, h**).

LIP activities within PD receptive field also showed this accumulation signature. VarCE grew with time and CorCE decayed with lag for perceptual decisions (**Extended Data Fig. 7c, d, 9c, f**), as expected for a genuine within-LIP accumulator. Because the signature is present for the perceptual decision but absent for the UR within the same LIP population, the UR deficit is a genuine dissociation, not a global failure of LIP to accumulate. This positive control reinforces the conclusions above.

VarCE and CorCE thus converge from two complementary angles. A late, externally injected signal both inflates the variance of late time bins, producing the late rise in UR-unit VarCE, and severs the temporal autocorrelation that a within-network accumulator would preserve. Both signatures are present in LIP and in the SUR agent, and both are absent from the IUR agent.

### Population geometry distinguishes local from remote implementation

The single-unit and VarCE analyses indicate that LIP UR-target activity behaves like an externally driven decision maker rather than a within-network evidence accumulator. Both analyses, however, require sorting individual units into UR- and PD-selective groups, and one might worry that the conclusion hinges on that categorization. We therefore asked whether the same architectural distinction is visible in the geometry of the population response, using all units without conditioning on single-unit labels.

The two architectures make opposite geometric predictions. Under the IUR architecture, PD and UR trials are generated by a single latent accumulator and should therefore occupy a common low-dimensional manifold, organized along a coherence axis and an RT axis. Under the UR architecture, UR trials should instead diverge from the PD manifold in a direction set by the external trigger, which should be largely independent of stimulus strength, yielding a UR subspace misaligned with the PD subspace. The misalignment should be greatest where the accumulator and the external signal diverge most.

To test this, we constructed low-dimensional manifolds in two complementary ways. Coh-PCA applies PCA to population activity binned by coherence and chosen direction, preserving the stimulus axis of the manifold (**Fig. 6a, b**). RT-PCA applies PCA to activity binned by RT and chosen direction, preserving the RT axis (**Fig. 6d, e**). For UR trials, the “chosen direction” is defined as the agent’s second-stage perceptual decision, so that PD and UR trials are matched on the final perceptual decision outcome. Any residual difference reflects the path to that outcome, not the outcome itself.

**Fig. 6.**
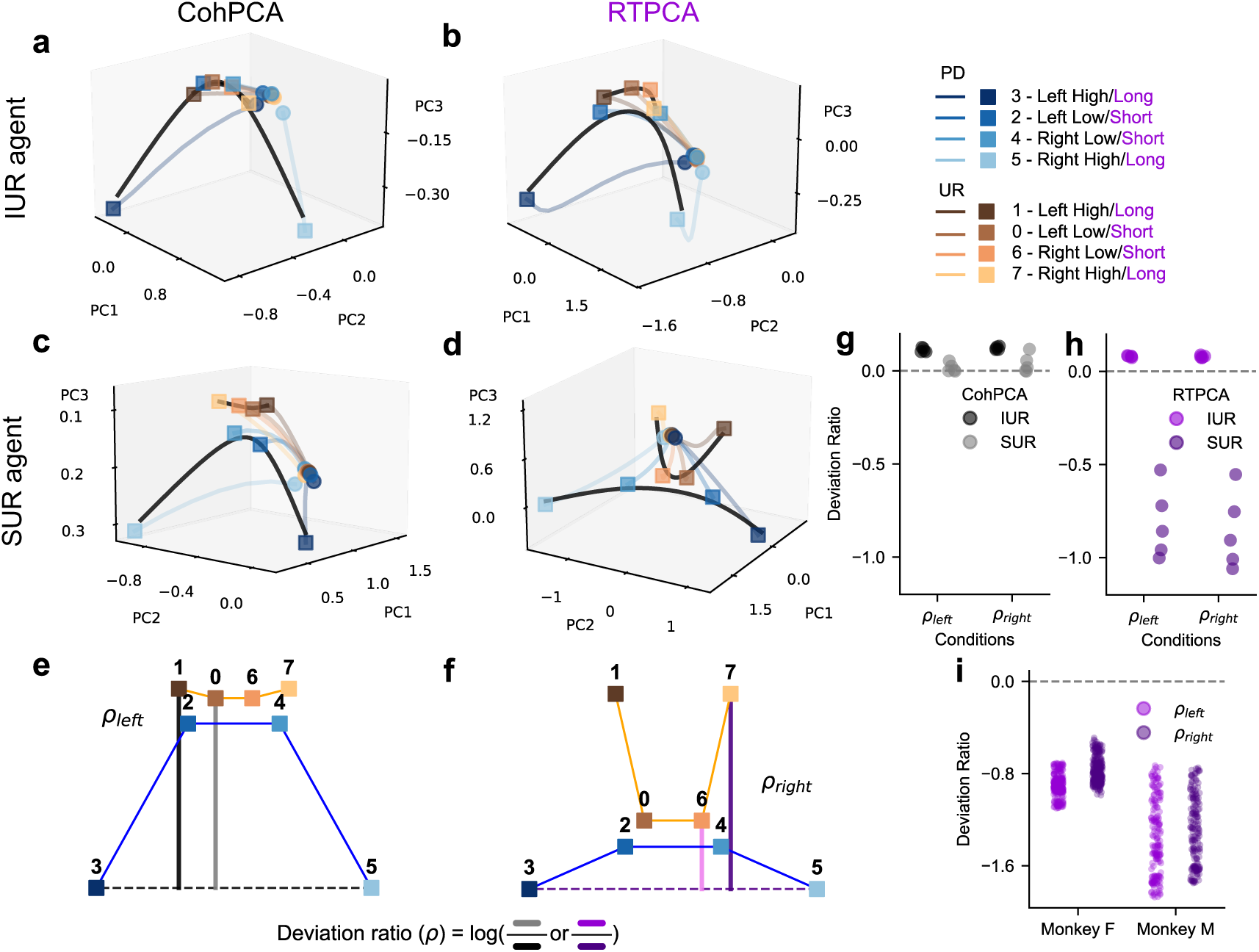
Population manifold geometry distinguishes integrated from segregated UR architectures through reaction-time PCA. **a-b.** Low-dimensional PCA manifolds from population activity binned by motion coherence and final chosen direction (Coh-PCA, **a**) or by reaction time and the final chosen direction (RT-PCA, **b**), using all units in the IUR agent (48 units, 30 sessions). Each trajectory is one combination of trial type (UR, warm colors; PD, cool colors), coherence (color shade), and chosen direction (left vs. right, indicated by brightness). Circles, stimulus onset; rectangles, 350 ms post-stimulus. Solid black curves connect terminal (rectangle) endpoints across conditions within a trial type with cubic-spline interpolation, defining the coherence-ordered terminal manifold. Dimmed colored curves connect each condition’s onset to its endpoint. For UR trials, chosen direction is the agent’s eventual directional report. **c-d.** Coh-PCA (**c**) and RT-PCA (**d**) on the SUR agent (48 units). **e-f**. Construction of the deviation ratio at the terminal time slice of the Coh-PCA (**e**) and RT-PCA (**f**) manifold. The black segment connects the two high-coherence PD endpoints (groups 3 and 5) as the reference line. For each UR point (points 0,1,6,7), we take its Euclidean distance to this line and compute the log ratio of the distance between the high- and low-coherence or long- and short-time as the deviation ratio. Ratios for the left and right points are denoted as ρ_*left*_and ρ_*right*_. **g–h.** Deviation ratios at successive post-stimulus time points for the Coh-PCA and RT-PCA. Windows span 0–350 ms post-stimulus for agents. The distinction between IUR and SUR agents under Coh-PCA is weak. **i.** Deviation ratios for the RT-PCA on monkey F (30 units), and monkey M (49 units), on *UR*_*in*_trials. Each point is one unit at one time point (1 ms). 200–700 ms (monkey F) or 200–583 ms (monkey M) for LIP after stimulus onset, truncated at median saccade latency to avoid contamination from motor signals. Results are robust to the window choice.

The two agents exhibit different geometries. In the IUR agent, UR and PD trajectories overlapped substantially in Coh-PCA (**Fig. 6a**), with UR trials clustering near the low-coherence PD choices, separated cleanly along an RT-ordered axis in RT-PCA (**Fig. 6d**). The Coh-PCA and RT-PCA manifolds were mutually consistent, indicating that PD and UR trials share a common representation. In the SUR agent, UR and PD trajectories diverged in both projections, with the largest divergence at high coherence under RT-PCA (**Fig. 6b, e**), where the externally triggered ramp is most clearly displaced from the PD manifold.

To quantify these differences, we defined a signed deviation ratio (Methods; illustrated in **Fig. 6c, f**). At the terminal time window, the line connecting the two high-coherence PD endpoints (**Fig. 6c, f**, groups 3 and 5) defines the PD axis. For each UR point (**Fig. 6c, f**, groups 0, 1, 6, and 7), we computed its Euclidean distance to this line and computed the log ratio of the distance between the high-(groups 0 and 1) and low-coherence (groups 6 and 7) UR groups (or, under RT-PCA, between long- and short-RT groups) as the deviation ratio. A positive index means the UR manifold’s curvature follows that of the PD manifold, while a negative index indicates a detachment from the PD manifold, the signature of a trajectory driven by an external trigger.

Under Coh-PCA, the indices are mostly positive in both agents and larger in IUR (**Fig. 6g, h**, black points), so coherence-based geometry cannot cleanly separate the models. Under RT-PCA, the two models diverge sharply: indices were predominantly positive in IUR (**Fig. 6g**, violet) but consistently negative in SUR (**Fig. 6h**, violet). The RT-PCA deviation ratio is thus the discriminating feature, and we applied it to the LIP data. Both monkeys’ LIP data replicated the signature of an external trigger in SUR. The RT-PCA deviation ratios were consistently negative on trials both in UR (**Fig. 6i**) and PD receptive field (**Extended Data Fig. 10a**).

These label-free population analyses converge with those from the single-unit and second-order results. LIP UR-target activity behaves like a late, externally injected signal occupying a population dimension deviant from the within-LIP PD accumulator and is inconsistent with a within-LIP accumulator architecture.

Together, these results suggest that the uncertainty estimation is unlikely to be a direct readout from the perceptual evidence represented in the LIP, and equally unlikely to be computed in a parallel uncertainty evidence accumulation process within the LIP.

## Discussion

We set out to answer two logically separable questions about how uncertainty is evaluated during an ongoing perceptual decision: whether the metacognitive signal is a readout of the perceptual decision variable or the product of a dedicated decision process, and, if the latter, whether that process is implemented within the circuitry that forms the perceptual choice or outside it. Behavioural modelling answered the first and the neural data, along with the RNN models, addressed the second. Together, the results suggest that concurrent metacognitive evaluation in this task is a parallel accumulation process implemented in distinct brain areas external to LIP.

This dissociation bears on how LIP’s role in decision making should be characterized. LIP carries evolving decision signals across a wide range of tasks, including perceptual judgments based on low-level visual features such as motion^32,33^, judgments based on higher-level visual features such as faces^38^, and probabilistic reasoning based on shapes^39,40^ or value^41,42^. One influential synthesis of these findings holds that LIP is generally engaged by any decision that links visual inputs to eye movements, irrespective of its cognitive content. Our results argue against this strong form of the hypothesis. The uncertainty response in our task was reported with a saccade and executed by the same effector as the perceptual choice, yet LIP activity preceding the two types of choices was qualitatively distinct. LIP ramping is therefore not a generic correlate of saccadic decision formation. This adds a novel within-task dissociation. The same neurons, recorded in the same sessions, with the same eventual motor output, show accumulator dynamics on perceptual decisions but not on uncertainty responses.

Our findings refine rather than contradict Kiani & Shadlen (2009)^4^, who reported that LIP firing rates predict whether the monkey will subsequently select a post-decision sure-bet target. The same logic applies to evidence that perturbing the sensory evidence perturbs choice and confidence together^29^. A shared sensory input is expected under both architectures we compare, and indeed both of our RNN agents are driven by a common evidence stream. Our task, in which uncertainty must be deliberated concurrently and can preempt the perceptual choice, places a stronger demand on the metacognitive process and reveals that LIP does not satisfy that demand. The LIP can serve as an input to an uncertainty computation, as Kiani & Shadlen (2009)^4^ demonstrated and as in our SUR model, but the computation that converts that input into a graded, bounded uncertainty signal appears to be carried out elsewhere.

A more direct point of comparison is the recent report by Vivar-Lazo & Fetsch (2025)^19^, who used a peri-decision wagering task and found that population activity in LIP carried concurrent and tightly time-locked signals for choice and wager. Although this result seems to contradict our finding, we regard the apparent disagreement as informative rather than contradictory: it may stem from differences in how metacognitive judgments are bound to motor reports, a pattern common in perceptual judgment^38,43–45^. In the peri-decision wagering paradigm, the choice saccade specifies a perceptual choice and a wager conjunctively. Because LIP neurons are spatially selective, the wagering signal in that task is, by construction, embedded in the same motor planning subspace that carries the perceptual decision. Consistent with this interpretation, Vivar-Lazo & Fetsch (2025)^19^ reported that the predictive coupling between choice strength and wagering probability holds only for contralateral choices, that is, only when the recorded population represents the winning perceptual accumulator and the corresponding chosen wager target. In our task, by contrast, the uncertainty response is reported by a saccade to a target that is not yoked to either perceptual alternative, so the uncertainty judgment can be expressed without commandeering the perceptual motor plan. Under these conditions, the segregation between perceptual evidence accumulation and uncertainty deliberation is exposed.

If this account is correct, then the two studies converge on a coherent picture. LIP implements the perceptual accumulator and represents whichever components of the metacognitive report can be expressed within its sensorimotor coordinate system, but it does not implement the underlying metacognitive computation. When confidence is bound to a saccade plan, that computation appears in LIP because its output is necessarily routed through the LIP motor representation. When confidence is decoupled from the perceptual saccade, as in our task, the absence of accumulator signatures for the uncertainty judgment makes clear that LIP is not the source of those signatures. This view also aligns with the behavioral demonstration by Kiani et al. (2014)^12^ that confidence is informed by both evidence and decision time, which suggests that the metacognitive variable depends on the same temporal dynamics that drive the perceptual decision but is computationally distinct from it. Concurrent does not imply colocalized, and the two are dissociable when the task allows.

A segregated architecture for concurrent metacognition has several appealing functional properties. First, decoupling the two processes allows the reliability of a decision to be estimated without that estimate being trivially determined by the decision itself. This is the property that allows opt-out and help-seeking behaviors to provide information beyond what is contained in the choice and reaction time, and it is consistent with the partial dissociation between accuracy and uncertainty responses observed behaviorally in our task and in others^8,36^. Second, a segregated metacognitive module in principle can be a domain-general “quality control” system that monitors multiple domain-specific lower-level decision circuits, rather than requiring each circuit to implement its own metacognitive machinery. Such an architecture is parsimonious, and it is broadly consistent with the proposal that metacognition is supported by a small number of higher-order regions that interface with many first-order circuits^25,26,46,47^. Third, a segregated module can integrate further information from experience and other cognitive factors that would be inappropriate to fold directly into a fast perceptual decision^9^.

If LIP does not produce the uncertainty judgment, where is it produced? A strong candidate is the pulvinar. Komura et al. (2013)^5^ found that pulvinar responses were reduced when monkeys selected an escape option, and, critically, that pulvinar inactivation increased escape choices without impairing categorization performance. The orbitofrontal cortex is also a likely candidate, given graded, modality-general confidence coding that predicts multiple confidence-guided behaviors^11,26,48^. Frontal candidates include the supplementary eye field^22^ and pregenual anterior cingulate and frontopolar cortex, whose reversible inactivation selectively impairs metacognitive judgments in macaques^28,49^ or humans^50^. Finally, the stereotyped, RT-independent presaccadic burst on UR trials resembles the bursting dynamics of the superior colliculus rather than the drift-diffusion dynamics of LIP^51^. These candidate regions are nonetheless drawn from escape, wagering, and confidence paradigms, none of which forcedly demands that the metacognitive judgment be committed to concurrently with the perceptual decision, as the present task does; testing them under this concurrent paradigm, for example by targeted inactivation, is the natural next step. If the UR signal reaching LIP is a commitment or threshold-crossing event rather than a developing decision variable, the abruptness of that rise is expected, and the accumulator that generated it should be sought upstream of the burst generator rather than in the structure that relays it.

Taken together, our results indicate that LIP, although long associated with perceptual evidence accumulation, does not accumulate the evidence underlying a concurrent uncertainty judgment, even when the two judgments share the same sensory inputs and motor outputs. Concurrent metacognition is therefore not a derivative readout of the perceptual decision circuit, but an independent process with its own anatomical substrate, which the present work leaves to be identified by future study.

## Methods

### Subjects and experimental procedures

Two adult male rhesus macaques (Macaca mulatta) were used in this experiment, with averaged body weights of 9.3 kg (subject F) and 6.8 kg (subject M), respectively, during the experiments. All experimental protocols were conducted in accordance with the guidelines approved by the Animal Care Committee of the Center for Excellence in Brain Science and Intelligence Technology, Chinese Academy of Sciences (protocol number CEBSIT-2021004).

#### Surgery and MRI

Prior to behavioral training, both monkeys underwent the implantation of a chronic titanium headpost. Following a two-month recovery period, the animals were trained to perform the behavioral task until they reached a satisfactory performance level. Structural Magnetic Resonance Imaging (MRI) scans were subsequently performed to accurately determine the location for the recording chamber. After the chamber implantation surgery, a manganese-enhanced MRI scan was conducted to confirm proper placement. All imaging procedures were conducted using a Siemens 3T scanner.

During the surgical procedures, sedation was initially induced with ketamine hydrochloride (10 mg/kg), followed by anesthesia induction and maintenance with isoflurane gas (1.5–2%, as required). Throughout the surgery, physiological parameters including body temperature, heart rate, blood oxygen saturation, and expired CO2 levels were continuously monitored.

#### Neural recordings

The implanted cylindrical acrylic recording chamber had an internal diameter of 19 mm. A plastic grid with 1 mm spacing was placed within the chamber to ensure accurate targeting of specific brain areas. Recordings were conducted using single tungsten electrodes (Alpha Omega), which were precisely maneuvered with the assistance of a Microdrive system (Electrode Positioning System, Alpha Omega). Neural data were recorded using the AlphaLab SnR System 2.0.4.5 (Alpha Omega) for high-fidelity acquisition of neural signals.

During each experimental session, an electrode was inserted into the ventral division of the lateral intraparietal area (LIPv), with precise localization guided by structural magnetic resonance imaging and identification of gray and white matter transitions during electrode penetrations^52^. Action potential waveforms were isolated in real-time using a window discriminator. Only neurons with clearly isolated waveforms were considered for recording and subsequent analysis.

We first mapped each neuron’s receptive field using a random-dots patch moved to various locations on the screen and then screened the neurons with the standard memory-guided delayed saccade task. Neurons were selected for further study only if their delay-period activity was significantly elevated above baseline for at least one target location and exhibited clear spatial selectivity.

#### Behavioral tasks and materials

The head-fixed monkeys were positioned in a primate chair, facing a 23.6-inch computer monitor located 70 cm from their eyes. The monitor was aligned such that its center corresponded with the midpoint between the subjects’ eyes. Behavioral tasks were executed using the MATLAB-based software MonkeyLogic. Eye positions were recorded using an infrared tracking system (EyeLink 1000) at a sampling rate of 500 Hz. A computer-controlled solenoid system was used to deliver the juice rewards.

#### Standard direction discrimination task

The monkeys participated in a reaction-time version of a random-dot motion direction discrimination task^33^. Each trial began when the monkey fixated on a central point and held for 500ms. Two choice targets were presented: one within the receptive field (RF) of the neuron under investigation and the other in the opposite visual hemifield. The choice targets were presented for a variable period before the onset of the motion stimulus, with this pre-stimulus interval randomly sampled (truncated exponential distribution, 500ms on average). This randomization discouraged anticipation of the stimulus onset and was crucial for training in the reaction-time task. After the pre-stimulus interval, a random-dot motion stimulus appeared within a 5°-diameter aperture centered at the fixation point, presented on a computer monitor with a frame rate of 60 Hz. Dots were drawn in three interleaved sets shown every third frame; on each update, each dot was displaced coherently along the motion direction with probability equal to the coherence, and otherwise replotted at a random position, yielding stochastic (geometrically distributed) dot lifetimes. The net direction of motion was toward one of the targets, with both motion direction and coherence randomly selected from the set {-25.6%, -12.8%, -6.4%, 0%, 6.4%, 12.8%, 25.6%} for each trial.

Once the motion stimulus was initiated, the monkey could indicate its choice at any time by executing a saccadic eye movement toward the corresponding choice target and holding the target for 200ms. The fixation point changed color at stimulus onset and reverted 200ms later, remaining so until the saccade. Correct choices were rewarded with a liquid incentive, and trials with 0% coherent motion were randomly rewarded. Incorrect responses resulted in a short time-out penalty (monkey F: 2000ms; monkey M: 750ms). Additionally, auditory feedback was provided to indicate an incorrect choice.

#### Concurrent uncertainty estimation task

Each trial began with the monkey fixating on a central point. After a 500ms delay, three targets were presented: two perceptual decision (PD) targets, identical to the standard direction discrimination targets, and one uncertainty response (UR) target, positioned orthogonally to the PD targets. Following a random delay, a random dot stimulus appeared. The monkey was allowed to indicate its choice by saccading to a choice target after the stimulus onset. At stimulus onset, the fixation point turned a new color. 200ms later, it returned to its original color until the saccade occurred. The period between stimulus onset and the choice, irrespective of the choice type (PD or UR), is referred to as the first stage.

If a PD target was chosen, the monkey was required to maintain fixation on it for 250ms. A reward was delivered for correct choices, while an incorrect choice resulted in a 750ms time-out penalty. Additionally, auditory feedback with different tones was provided to indicate whether the choice was correct or incorrect. If the monkey selected the UR target, a 25.6% coherence random dot stimulus, with the same motion direction as the first stimulus, was presented. As the cost for choosing the UR target, there was a random delay (truncated normal distribution. Monkey F: 400ms; Monkey M: 800ms on average) before the onset of the second motion stimulus. The monkey could, then, choose a PD target at any point by making a saccade and holding the fixation for 250ms. A correct choice resulted in a reward equal to that of first-stage perceptual decisions, while an error was signaled with auditory feedback, followed by a time-out penalty (monkey F: 2000ms; monkey M: 750ms). Likewise, the interval from the second stimulus onset to the choice is defined as the second stage.

### Behavioral data analysis

All behavioral analyses were conducted based on the monkeys’ performance during recording sessions. Trials exhibiting noticeable and sustained primate chair shaking were excluded, along with those where the trial was terminated before a decision was made. We applied a logistic regression to fit the proportion of rightward choices to derive psychometric curves.

#### Drift diffusion models

We used two drift diffusion models^53^ to fit the behavior. The first model, termed the single-accumulator model, places the UR boundary inside the PD evidence space. The drift diffusion process is described as

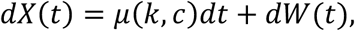

where *X*(*t*) represents the decision variable. The drift term is given by *μ*(*k*, *c*) = *k c* and *W*(*t*) is the standard Wiener process. The boundaries change linearly. For PD, the correct and wrong boundary are *b_PD_* − *τ_PD_t* and −*b_PD_* + *τ_UR_t*, respectively. The UR boundaries consist of two segments: *τ_UR_*(*t* − *t_peak_*) when *X*(*t*) ≥ 0 and *t* > *t_peak_*, and −*τ_UR_*(*t* − *t_peak_*) when *X*(*t*) < 0 and *t* > *t_peak_*. They form a triangular inner UR region as illustrated in **Fig. 2a**.

The second model, termed the parallel-accumulator model, treats the UR process as a separate single-boundary accumulator, competing with the perceptual decision process, which was modelled as an evidence accumulator for correct and incorrect responses. We represent their joint dynamics using a system of coupled drift-diffusion equations in a matrix form:

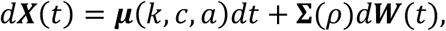

where ***X***(*t*) = (*X_PD_*(*t*), *X_UR_*(*t*))′ are the PD and UR evidence accumulation processes, respectively. The drift vector is ***μ***(*k*, *c*, *a*) = (*k c*, *a* − *k c*)′, and the dif fusion matrix 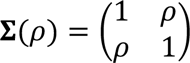. ***W***(*t*) denotes a two-dimensional standard Wiener process. Within this framework, the PD boundaries are defined as, *b_PD_* − *τ_PD_t* and −*b_PD_* + *τ_PD_t*, while the UR boundary is given by *b_UR_* − *τ_UR_t*. For computational efficiency, a secondary absorbing boundary is placed sufficiently far from the primary UR boundary.

We employ Markov-based methods^54,55^ to approximate the reaction time distributions, which subsequently allow for the derivation of accuracy and UR proportions. Both models incorporate a non-decision time component, modeled as a normal distribution with independently fitted mean and variance parameters. In total, the single-accumulator model comprises 7 parameters, while the parallel-accumulator model contains 10. For model fitting, we optimize both reaction time likelihoods and weighted mean squared errors using the Nelder-Mead algorithm^56^ in *scipy*, evaluating multiple initial conditions to identify the global optimum. Model comparison used the Bayesian Information Criterion (BIC) on the pointwise reaction-time likelihoods (**Fig. 2c**). The wMSE was optimized under a Gaussian-error model, so that *BIC* = *k ln N* − 2 *ln L*, with the wMSE corresponding to −*ln L* up to an additive constant. The number of free parameters was *k* = 7 for the single-accumulator model and *k* = 10 for the parallel-accumulator model. The number of observations was *N* = 16: four quantities (accuracy, mean RT on PD trials, mean RT on UR trials, and UR proportion) at each of the four coherence levels. (**Fig. 2c, d**).

### Neural data analysis

All neurons that met the selection criteria were included in the subsequent data analysis. In the memory-delay-saccade task, when the target appeared within the neuron’s receptive field, the neuron exhibited an increased response to target onset or the saccade. For several analyses, units with fewer than 600 trials were additionally excluded to ensure statistical reliability. Additionally, the neuron maintained an elevated firing rate throughout the delay period.

#### PSTH

Neurons were categorized based on condition, receptive field, and trial type. Conditions included the standard (STD) and concurrent uncertainty estimation (CUE) tasks, while receptive fields (RF) were categorized as the perceptual decision (PD) targets (*T_in_* and *T_ou_*_*t*_) or the uncertainty response (UR) target. Trial types were defined as either directly choosing PD or initially selecting UR. Trials were further grouped by coherence/reaction time and motion direction. Motion direction was not distinguished in the UR trials.

#### K-nearest neighbor decoder

To decode PD versus UR choices, a *K*-nearest neighbors (KNN) classifier is employed with *scikit-learn*. This nonparametric method is chosen because it does not assume linear separability and can capture multimodal class boundaries that may arise across heterogeneous stimulus conditions. It assigns class labels based on the majority vote among the *K* most similar training samples in feature space. Selection of *K* involves a bias–variance trade-off: small *K* increases sensitivity to noise, whereas large *K* may oversmooth decision boundaries and introduce bias. A value of *K* = 5 is adopted, consistent with common practice (brute-force Euclidean distance) for binary classification tasks with stratified 5-fold cross-validation. Decoding accuracy was computed as the mean fraction of correctly classified test trials across folds. As a chance-level control, the same procedure was repeated with training labels randomly permuted within each fold (test labels unchanged), using an identical fold structure.

To assess significance, decoding accuracies (real vs. shuffled) were compared across neurons using a one-sided Wilcoxon signed-rank test, separately for each time point (**Fig. 4g, h**) or time window (**Fig. 4i, j**).

#### Variance Analyses

Following methods in the previous study^37^, we quantify characteristic variance and autocorrelation features for individual neurons with Variance of the Conditional Expectation (VarCE) and Correlation of Conditional Expectation (CorCE). For LIP data, only trials in which the final choice aligns with the neuron’s preferred direction are included. For RNN units, only trials with the final choice aligned to the PD or UR directions, specifically, leftward choices are retained. Reorienting the selected direction across datasets does not alter the results qualitatively.

For each unit and each 100ms time bin, VarCE is defined as,

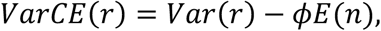

where *r* is a random variable capturing the unit’s spike count (LIP) or mean firing rate (IUR and SUR); *E*(*n*) is the corresponding mean spike count or mean firing rate; *φ* is the Fano factor, assumed constant across time bins for a given unit. The term *φE*(*n*) corresponds to the point process variance, which scales linearly with the mean and reflects the intrinsic variability of the spiking process under a stationary Poisson-like assumption.

Similarly, the CorCE between time points *i* and *j* is expressed as

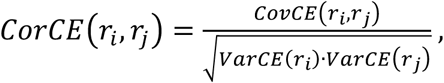

where *i* and *j* are time points. Under the assumption of temporal independence across time bins for the underlying point process. Consequently, the conditional

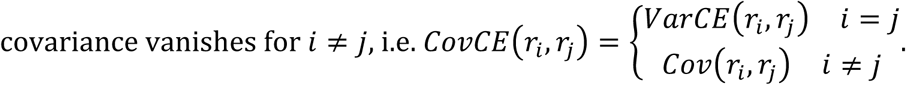

#### Principal component analysis

To ensure comparability between single-neuron electrophysiological data and model outputs, despite the latter generating population activity simultaneously across units, the units are treated as independent. Our analyses confirmed that neither population-level PCA nor the inclusion of unit activities recorded from distinct sessions affects the primary conclusions.

Let the data tensor D ∈ ℝ*^N^*^×*T*×*M*^, where *N* denotes the number of units, *T* the duration in time bins of the longest trial within the epoch of interest, and *M* the total number of trials. Trials are averaged separately for each unit across conditions defined either by choice and motion coherence (Coh-PCA) or by reaction time quartiles (RT-PCA). Only activities spanning the interval from stimulus onset to the termination of the first PD and UR are included.

The PSTH for each unit is aggregated as a PSTH data matrix D*_pca_* ∈ ℝ*^CT^*^×*N*^, where *C* is the number of behavioral conditions. Each unit’s PSTH is detrended prior to PCA to remove slow drifts. The resulting principal components are projected into a low-dimensional space of dimension 3, yielding a transformed tensor in _ℝ_*C*×*N*×*d*.

### RNN models

#### Reinforcement learning

The integrated and segregated UR agents are trained using proximal policy optimization (PPO)^57^, employing a framework adapted from previous work^58^. Training data are generated online at each time point through interaction with a predefined task environment. Model parameters are updated after every 20 trials using online gradient-based optimization. Upon completion of training, the model can generate new data via environment interaction without requiring external feedback.

To minimize the value function estimation error while maximizing expected cumulative reward and encouraging exploration, we minimize the following target function:

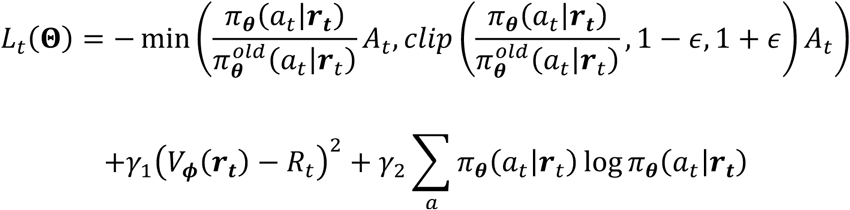

where **Θ** = {***θ***, ***φ***, ***J***, ***B***, *c*} denotes the set of learnable parameters in the policy *π**_θ_***, value function *V****_φ_***, recurrent weights ***J***, input weights ***B***, and recurrent bias *c*; 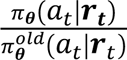 is the probability ratio between the current and previous policies, where ***r***_*t*_ is the vector of units firing rates from a vanilla recurrent neural network discussed later. *ε* = 0.3 is the clipping parameter^59^. *γ*_1_ = 0.5 and *γ*_2_ = 0.01 are hyperparameters governing value and entropy regularization, respectively. The final term corresponds to the entropy of the policy distribution, which serves to encourage exploration.

The value function *V****_φ_***(***r***_*t*_) is a linear readout of ***r***_*t*_ with an added bias term. The stochastic policy *π**_θ_***(*a*_*t*_|***r***_*t*_) is implemented as a softmax distribution over actions, parameterized by a linear transformation of ***r***_*t*_. Actions are sampled according to this distribution and executed in the simulation environment. The return *R*_*t*_ is defined as the discounted sum of future rewards:

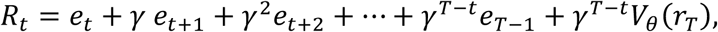

With temporal discount factor *γ* = 0.99. The advantage estimate is given by *A*_*t*_ = −*V_θ_*(*r*_*t*_) + *R*_*t*_, where *e*_*t*_ denotes the reward supplied by the environment at time *t*.

#### RNN structure

Both IUR and SUR agents incorporate an RNN comprising 48 recurrent units. Their dynamics are governed by the differential equation,

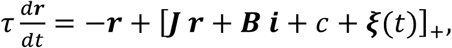

where ***J*** and ***B*** are recurrent and input weight matrices, respectively, *c* is a constant bias; and ***ξ***(*t*) is Gaussian white noise with zero mean and covariance *Cov*e***ξ***(*t*)*_i_*, ***ξ***(*t*)_j_f = 2*τ σ*^2^ *δ*(*i* − *j*)/ Δ*t*, approximated during simulation with *σ* = 0.1. The input vector ***i*** comprises four channels for IUR and five for SUR (the fifth channel conveying the external UR signal). Temporal discretization is performed with Δ*t* = 10*ms*, corresponding to 50*ms* in simulated task time.

#### Temporal structure

For IUR, each trial starts with the inter-trial interval (ITI) period, followed by the fixation period and then the stimulus period. If the agent selects the UR action during the stimulus period, the environment transitions to the UR-chosen period. The temporal structure of the whole trial determines the input channels ***i***_*t*_ presented to the reinforcement learning agent at time *t*.

#### Input channels

IUR uses four input channels, while SUR includes an additional one. The fixation channel is set to 1 during the ITI period and 0 otherwise. The UR channel is set to 0 during the second stage period and 1 otherwise, encoding task progression. There are two stimulus channels: 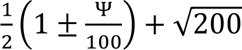· 0.01*N*(0,1) during the stimulus period, where Ψ is the coherence level with a moving direction chosen from a randomly signed 2,4,6,8,13,26,52,78 with equal probabilities^60^. The simulation employs this 8-level coherence grid in place of the exact behavioral set, since the model is capable of achieving high accuracy by the end of training but converges slowly in its early stages. During the UR-chosen period, Ψ is set to ±78, which is the highest coherence. SUR additionally includes an external UR trigger channel, set to 1 when the external UR module produces a UR decision and 0 otherwise.

#### Action channels

Four actions are available: fixation, left, right, and UR.

#### Performance-contingent feedback

During the ITI period, all actions are permissible; during fixation, only fixation is allowed; during stimulus/UR-chosen, selecting the PD option terminates the trial and yields reward or punishment based on response accuracy. UR selections incur a switching cost, with reward magnitudes adaptively adjusted during training to shape behavior. This fixed penalty serves as a functional stand-in for the behavioral time cost for convenience, that is, the delay before the second-stage stimulus, which the model absorbs into the reward schedule instead of simulating it as explicit elapsed time. For SUR, the agent must execute UR within 200ms (4 time points) following the receipt of the external UR signal via the eUR channel. Failure results in trial termination and a penalty.

#### Training curriculum

The training proceeds in three stages. (i) initial training without incentive for UR selection, using high-coherence stimuli (large |Ψ|), establishes the agent’s acquisition of trial-structure dynamics and the mapping between stimulus features and response accuracy; (ii) subsequent introduction of low-coherence stimuli (small |Ψ|) with reward for correct decisions following UR invocation enables the agent to learn to use UR; (iii) gradual reduction of UR-reward magnitude, combined with imposition of a fixed switching cost, shapes UR strategies. The training protocol for SUR mirrors that of IUR, with UR execution timing added as an additional constraint.

We implemented task-specific adaptations to align simulated behavior with empirical subject performance. The training pipeline follows a sequential curriculum: each stage must meet a predefined performance criterion before progression to the next, analogous to controlled behavioral training in nonhuman primates but with quantifiable decision thresholds.

#### Simulations

We generated 30 sessions, each consisting of 10,000 trials, for both IUR and SUR agents. Trials with premature termination (“break trials”) are excluded (break rates: IUR = 4.48 ± 0.22%; SUR = 5.85 ± 0.20%; SUR without external UR signal = 2.59 ± 0.16%). Analyses of behavioral and neuronal dynamics used the first 8 sessions per dataset; PCA analysis employed the full 30-session set.

#### Units choice probability

The policy function is defined as ***π***(***r***) = *softmax*(***A r*** + ***b***), where ***π***(***r***) denotes the probability distribution over *C* possible actions (fixation, left, right, UR). For each time step and trial, the Jacobian of *π* with respect to the activity vector ***r*** is given by

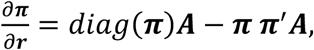

where 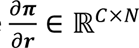. *N* is the number of units. For each unit and action, we compute the mean Jacobian element across all trials and time points, separately for positive 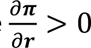 and negative 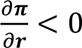 contributions. This decomposition identifies units with activities correlating with action probability. Additionally, we report the ensemble-averaged Jacobian across all trials and time points to characterize the net directional sensitivity of each unit to activity perturbations (**Extended Data Fig. 11**).

## Data and code availability

Data and code are available from the corresponding author upon reasonable request.

## Acknowledgments

We thank Roozbeh Kiani and Michael Shadlen for insightful discussions during the study, and Rui Zhu, Wei Kong, and Lu Yu for their help in all phases of the study. The authors declare no competing financial or nonfinancial interests.

## Author contributions

T.Y. designed the study. X.L. and Y.C. performed the experiments and collected the data with assistance from R.S. X.L. designed the RNN models. X.L., Y.C., and T.Y. analyzed the data and wrote the paper.

## Extended Data figures

**Extended Data Fig. 1.**
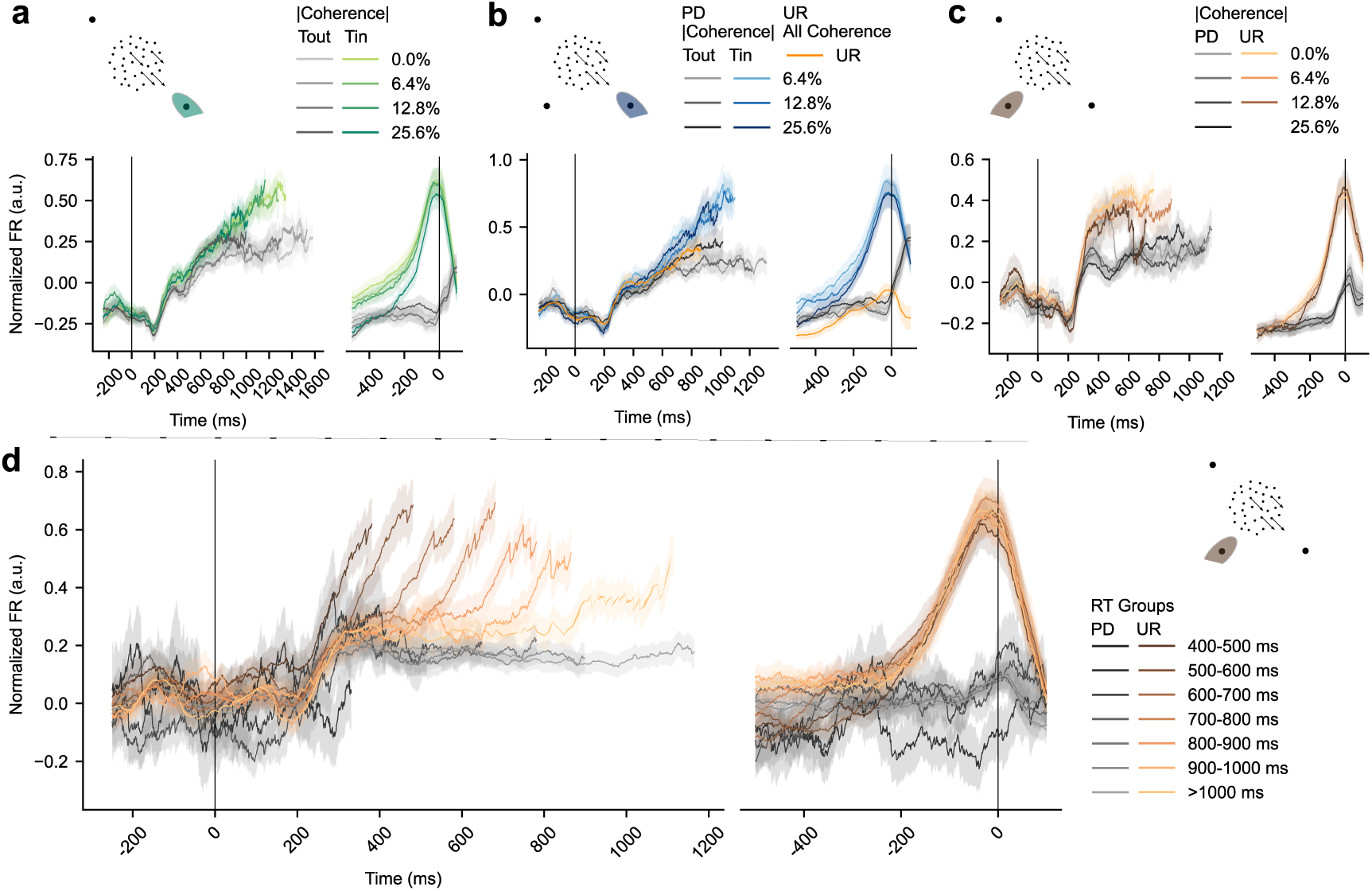
LIP responses across receptive-field configurations and tasks, monkey. **F.** Following the same labels as Fig. 3.

**Extended Data Fig. 2.**
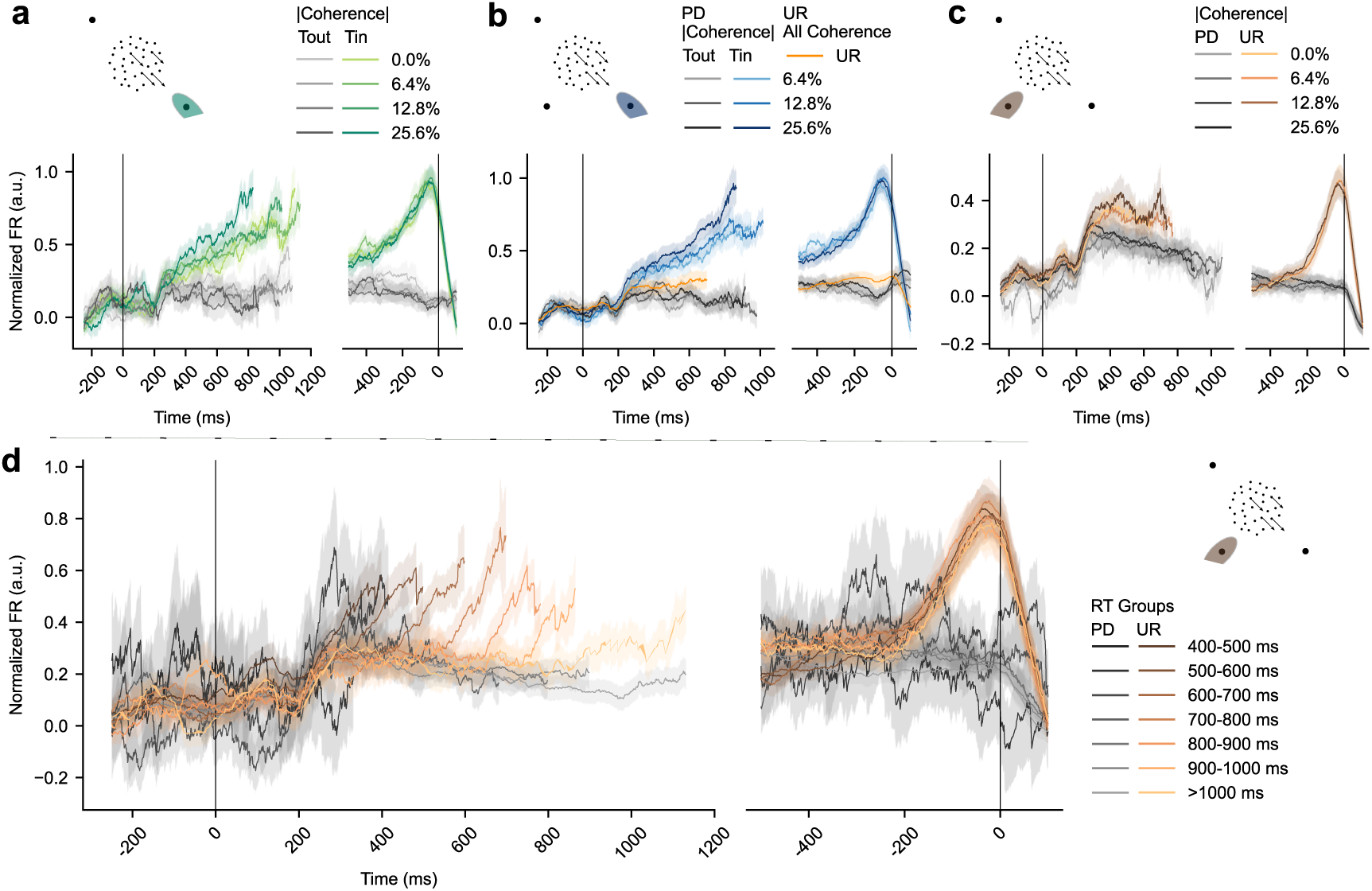
LIP responses across receptive-field configurations and tasks, monkey. **M.** Following the same labels as Fig. 3.

**Extended Data Fig. 3.**
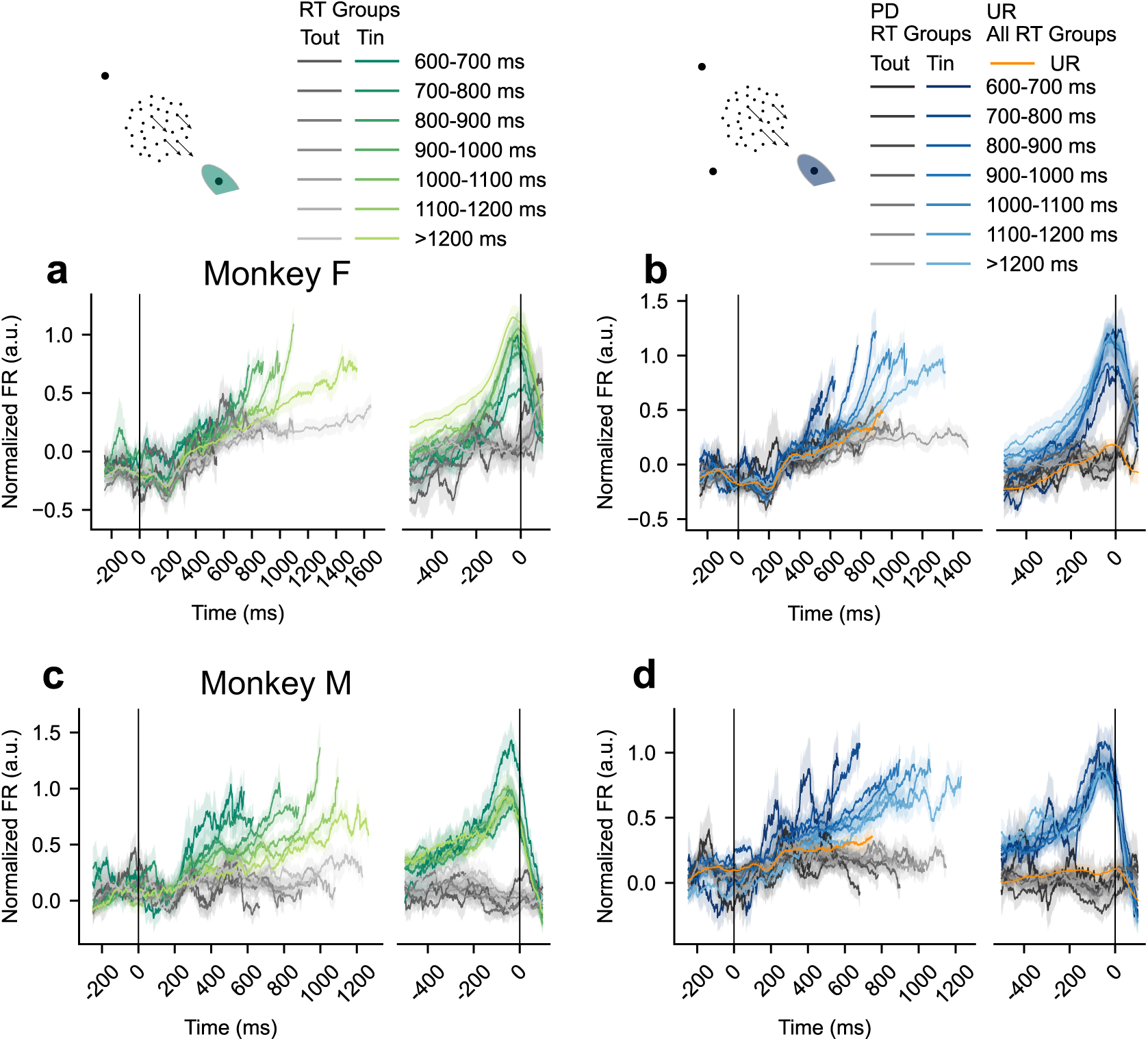
LIP responses in the STD and CUE tasks, with the UR target in the response field, grouped by reaction time. Conditions with too few trials (**a**, <600ms in STD, *T_in_* 47 trials in 11 units, *T_ou_*_*t*_ 48 trials in 11 units; **b**, <600ms in PD, *T_in_* 86 trials in 20 units, *T_ou_*_*t*_ 45 trials in 12 units; **c**, <600ms in STD, *T_in_* 16 trials in 10 units, *T_ou_*_*t*_ 42 trials in 20 units; **d**, <600ms in PD, *T_in_* 28 trials in 11 units, *T_ou_*_*t*_ 33 trials in 22 units) are not plotted here. For the motion-onset-aligned traces, each condition was truncated at the across-unit average of the single-unit median RTs. The UR condition in **b, d** was truncated at three-quarters of that value, because UR reaction times were substantially shorter than PD reaction times. Colored lines except orange in **b, d**: *T_in_* trials; Grey lines: *T_ou_*_*t*_ trials. Individual monkeys showed similar patterns.

**Extended Data Fig. 4.**
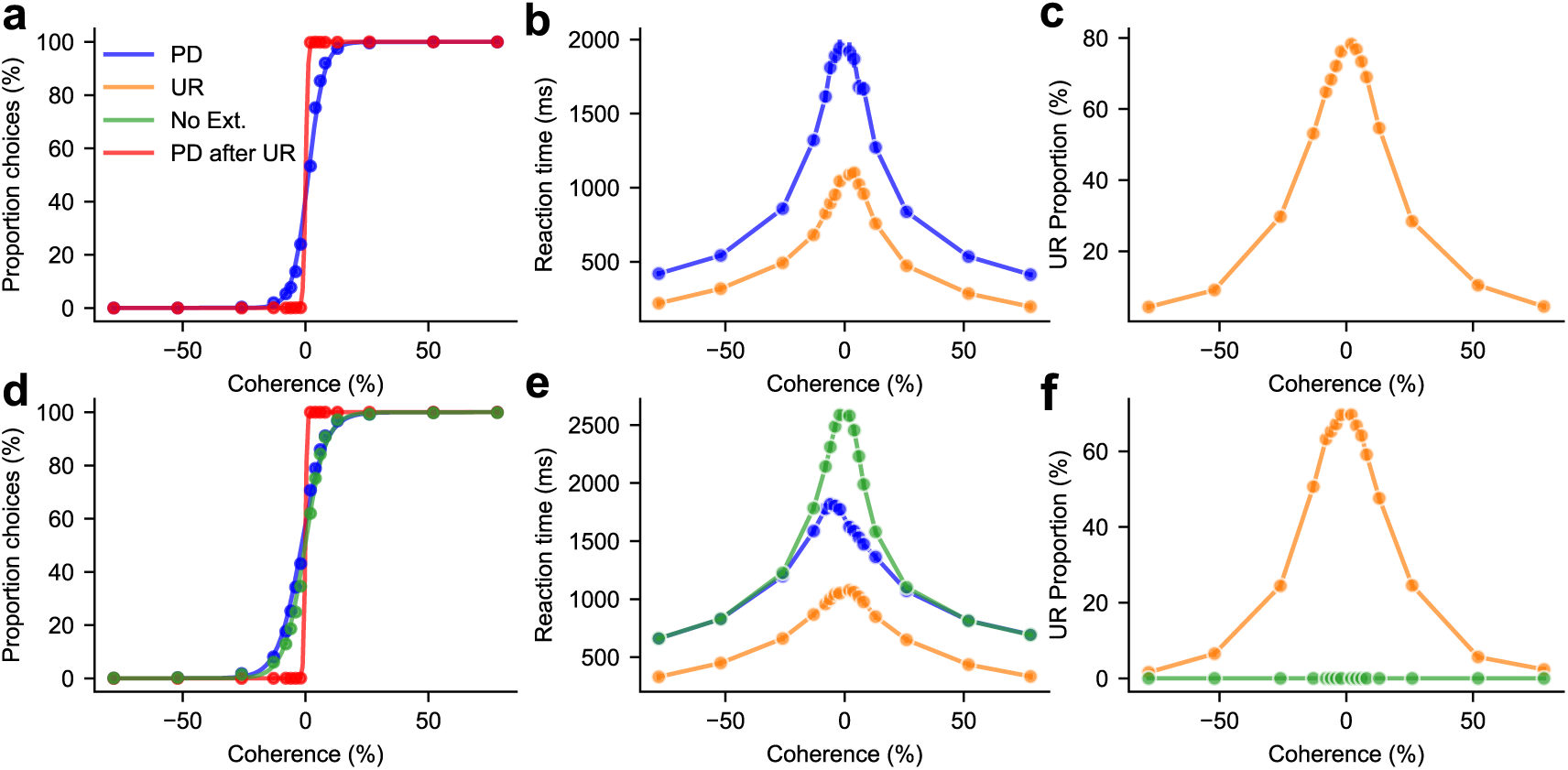
Behavioral metrics from IUR and SUR agents. **a-c** and **g-i**, Psychometric curve, reaction time and UR proportion as a function of coherence levels (IUR, *n_sess_* = 8; SUR, *n_sess_* = 8). **d-f** and **j-l**, Correct rate, reaction time, UR proportion across RT quintiles for the same IUR and SUR datasets, respectively.

**Extended Data Fig. 5a.**
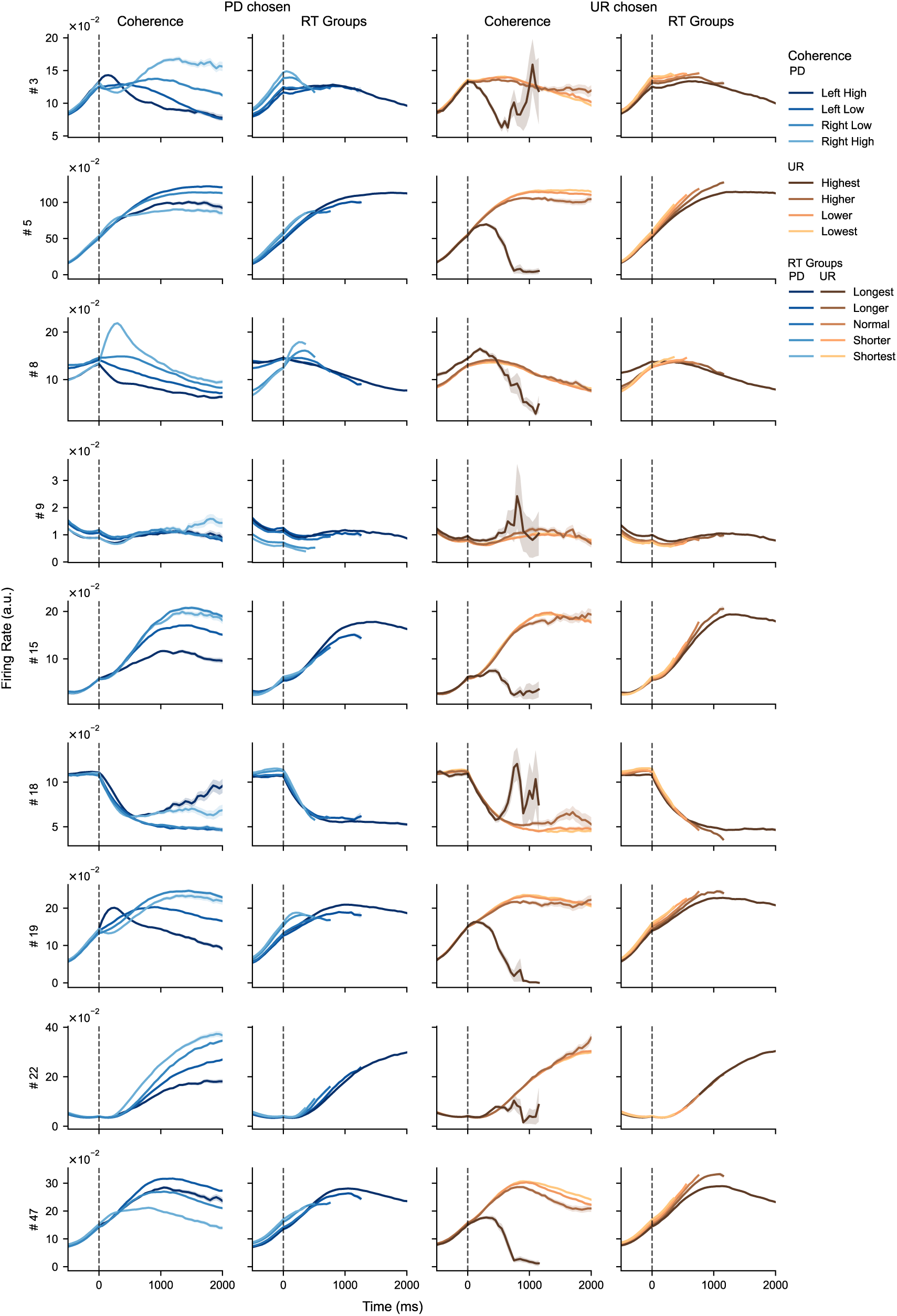
Normalized firing rates of positive UR units in the IUR agent. Dashed vertical lines indicate the stimulus onset. The last column, UR-chosen grouped by RT quintile, does not exhibit the same temporal pattern observed in the empirical neuronal data.

**Extended Data Fig. 5b.**
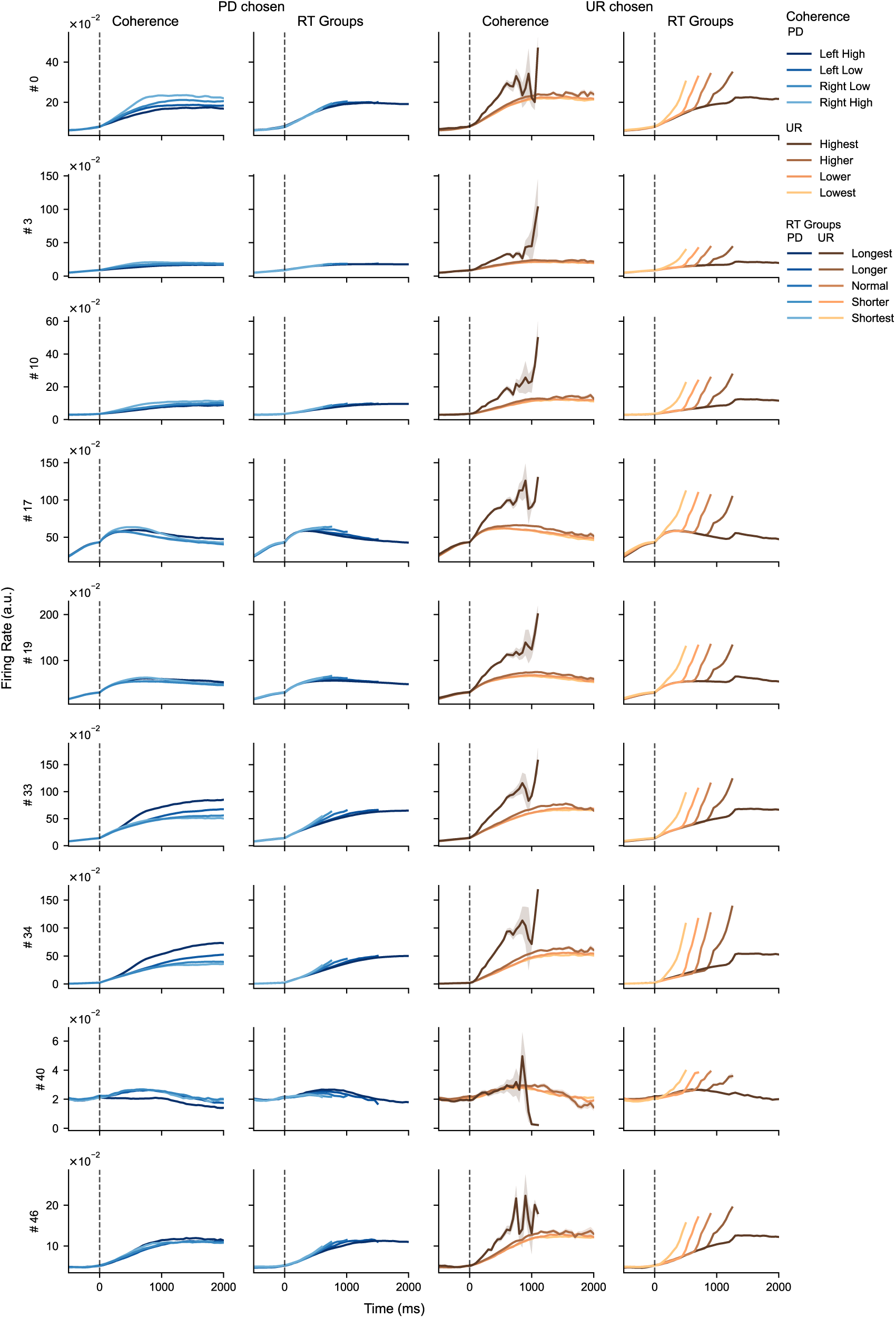
Normalized firing rates of positive UR units in the SUR agent. Dashed vertical lines indicate the stimulus onset. The last column, UR-chosen grouped by RT quintile, exhibits the same temporal pattern observed in the empirical neuronal data.

**Extended Data Fig. 6.**
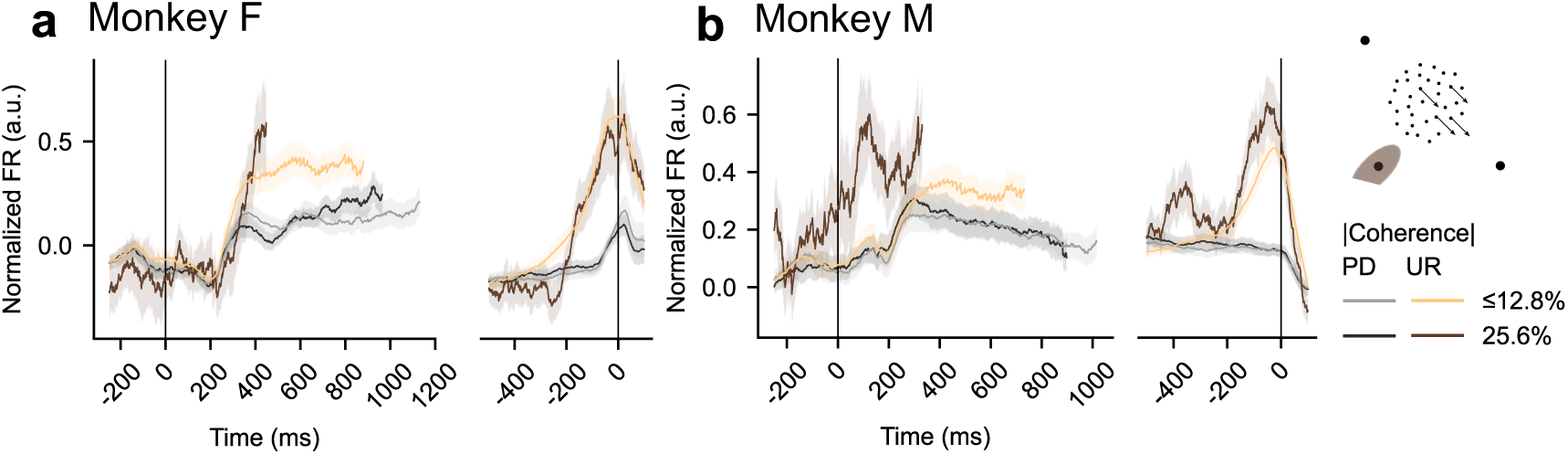
CUE-task LIP responses (UR target in the response field) grouped by coherence, including 25.6% coherence.

**Extended Data Fig.7.**
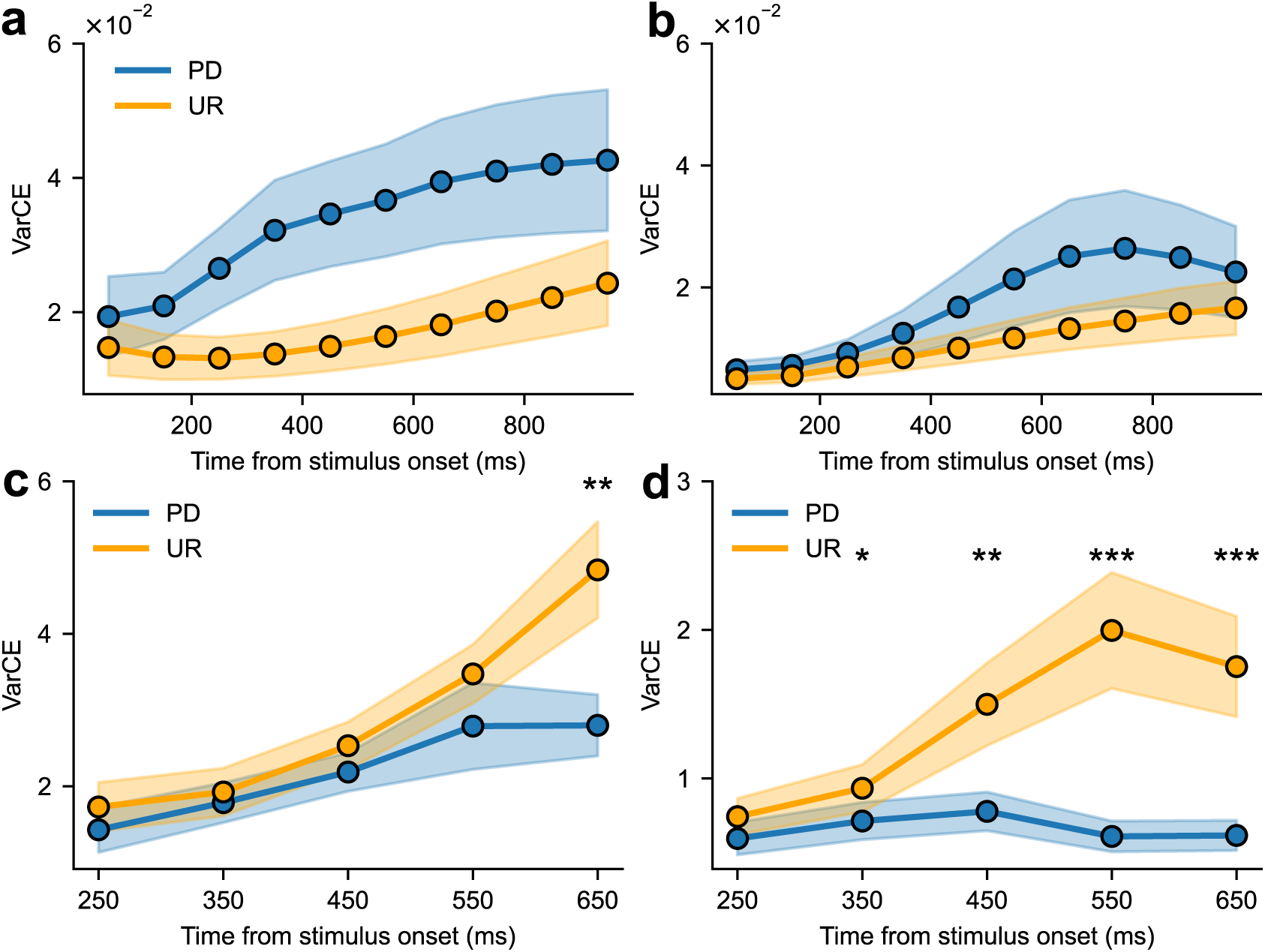
VarCE estimates. **a-b**, all units across IUR and SUR agents, respectively. **c-d**, units choosing PD in PD receptive field for monkey F and M, respectively.

**Extended Data Fig. 8.**
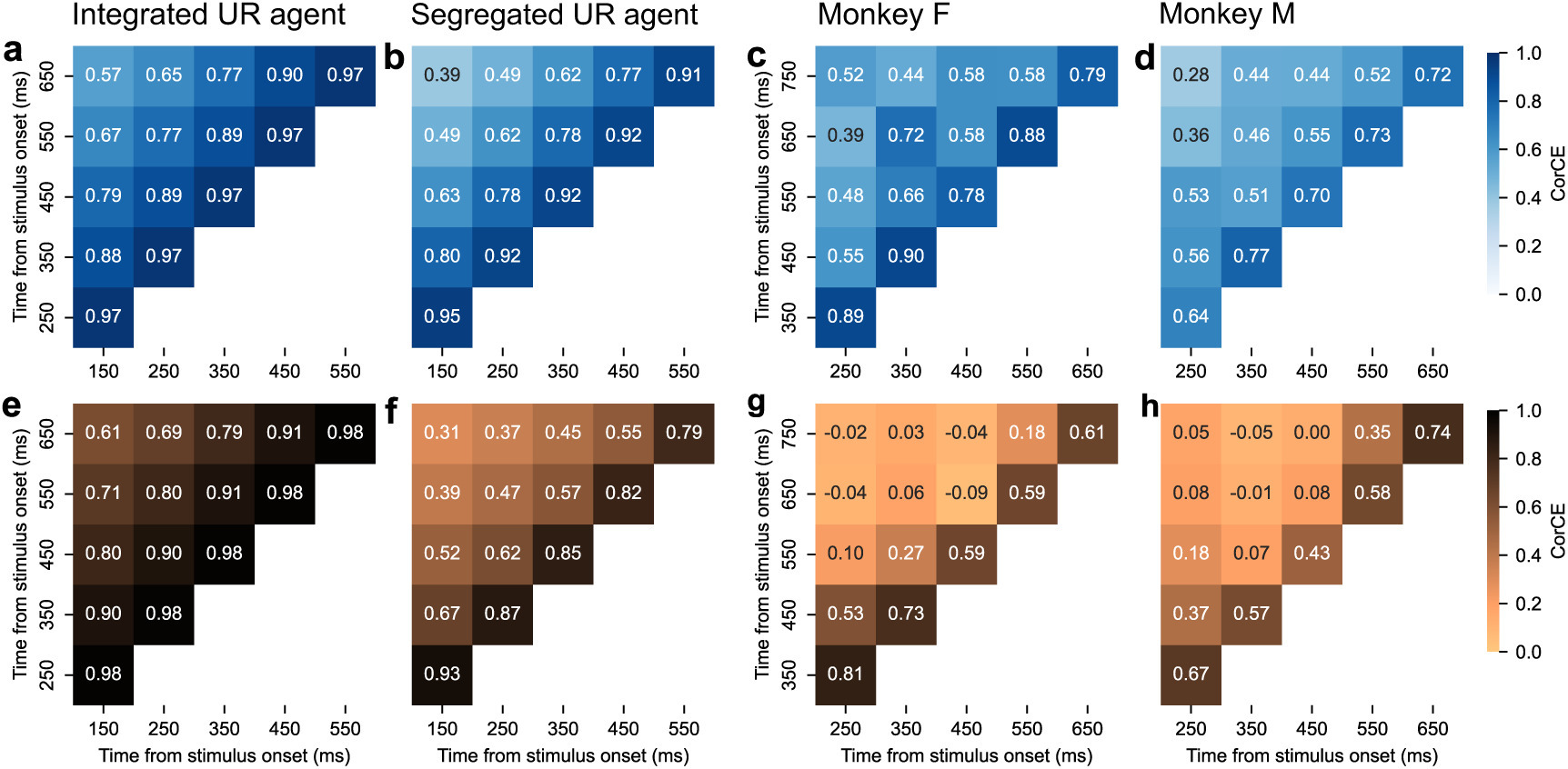
CorCE estimates for choosing PD and UR. **a-b**, CorCE for PD and UR in the IUR agent, respectively. **c-d**, corresponding results in the SUR agent. **e-f** and **g-h**, CorCE for UR choices within the UR receptive field for Monkey F and Monkey M, respectively.

**Extended Data Fig. 9.**
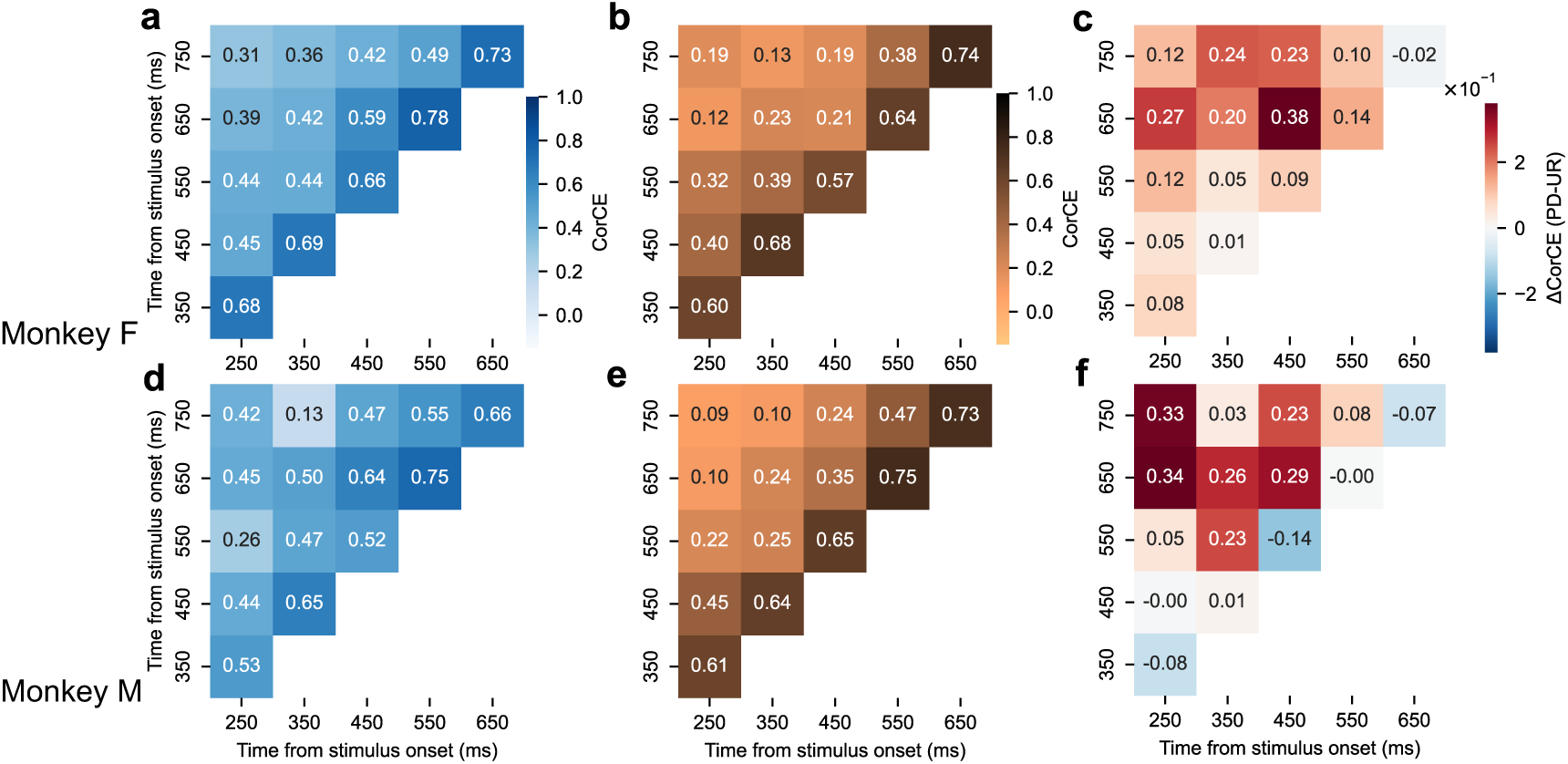
CorCE estimates for choosing PD and UR in LIP data, along with their differential contributions within the PD receptive field. **a-c**, Monkey F. **d-f**, Monkey M.

**Extended Data Fig. 10.**
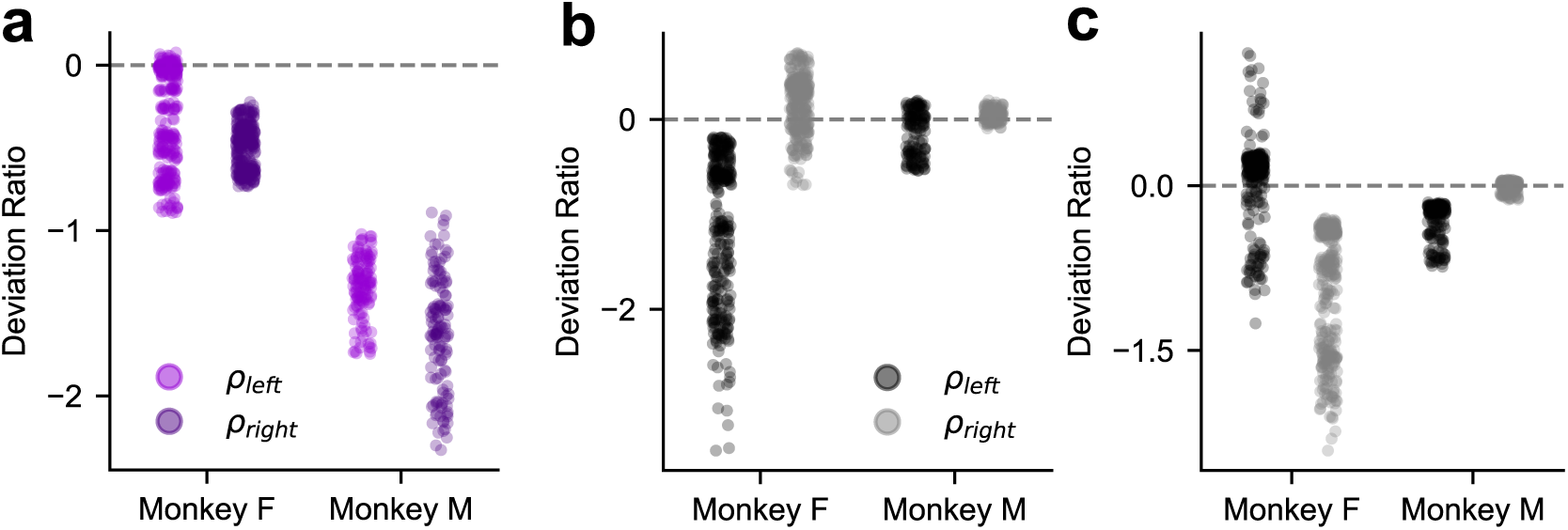
Deviation ratios at successive post-stimulus time points for the Coh-PCA and RT-PCA. **a.** RT-PCA in PD receptive field. For UR receptive field, see Fig. 6i. **b, c.** Coh-PCA on PD and UR receptive field, respectively.

**Extended Data Fig. 11a.**
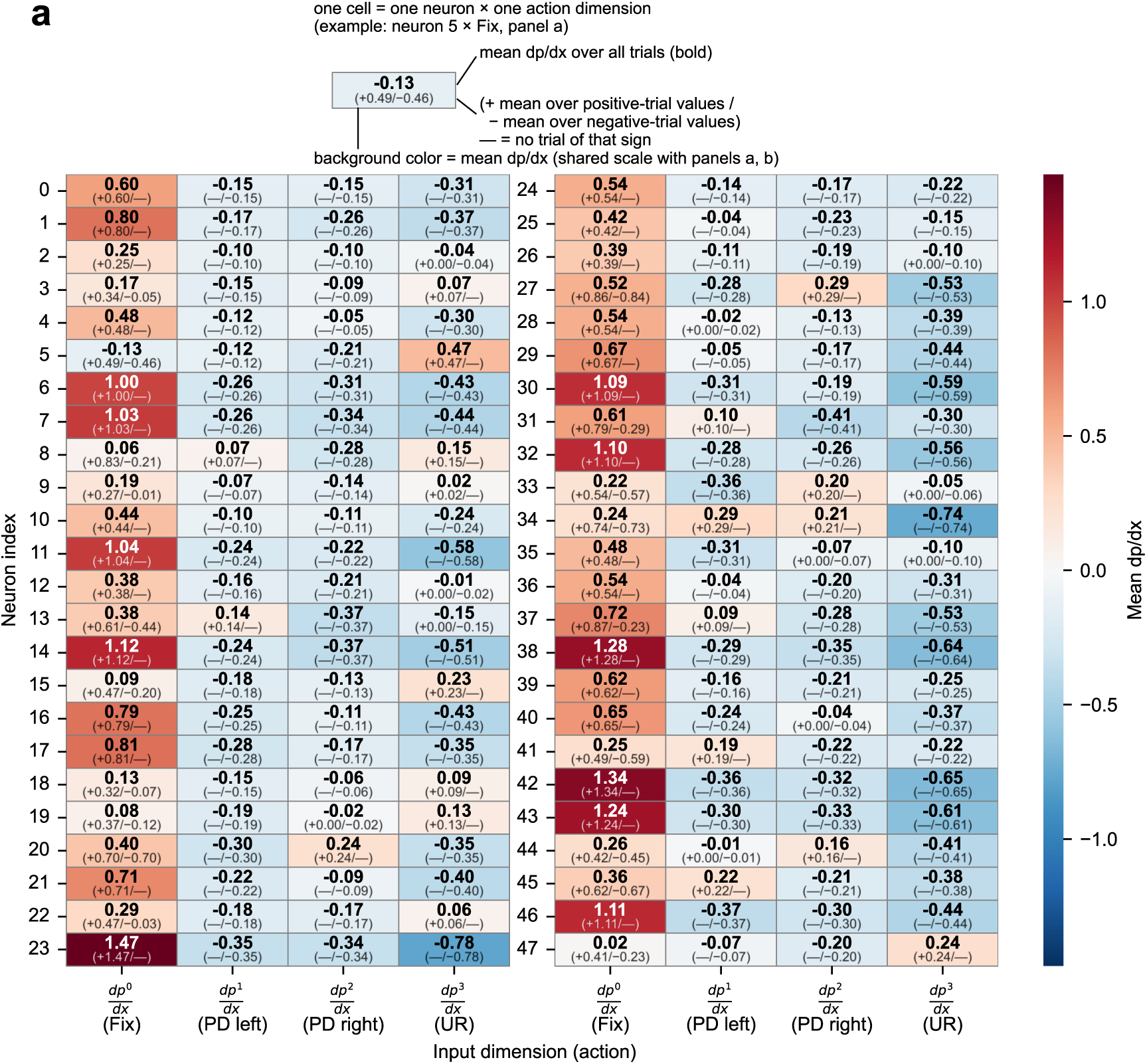
Choice probability contributions for individual units in the IUR agent. Each block reports, for a given unit, the mean overall contribution to the corresponding choice probability; values in parentheses denote mean contributions conditional on the sign of the effect (positive or negative). A dash indicates no contribution. Positive UR units are defined as those exhibiting exclusively positive contributions to the IUR choice probability.

**Extended Data Fig. 11b.**
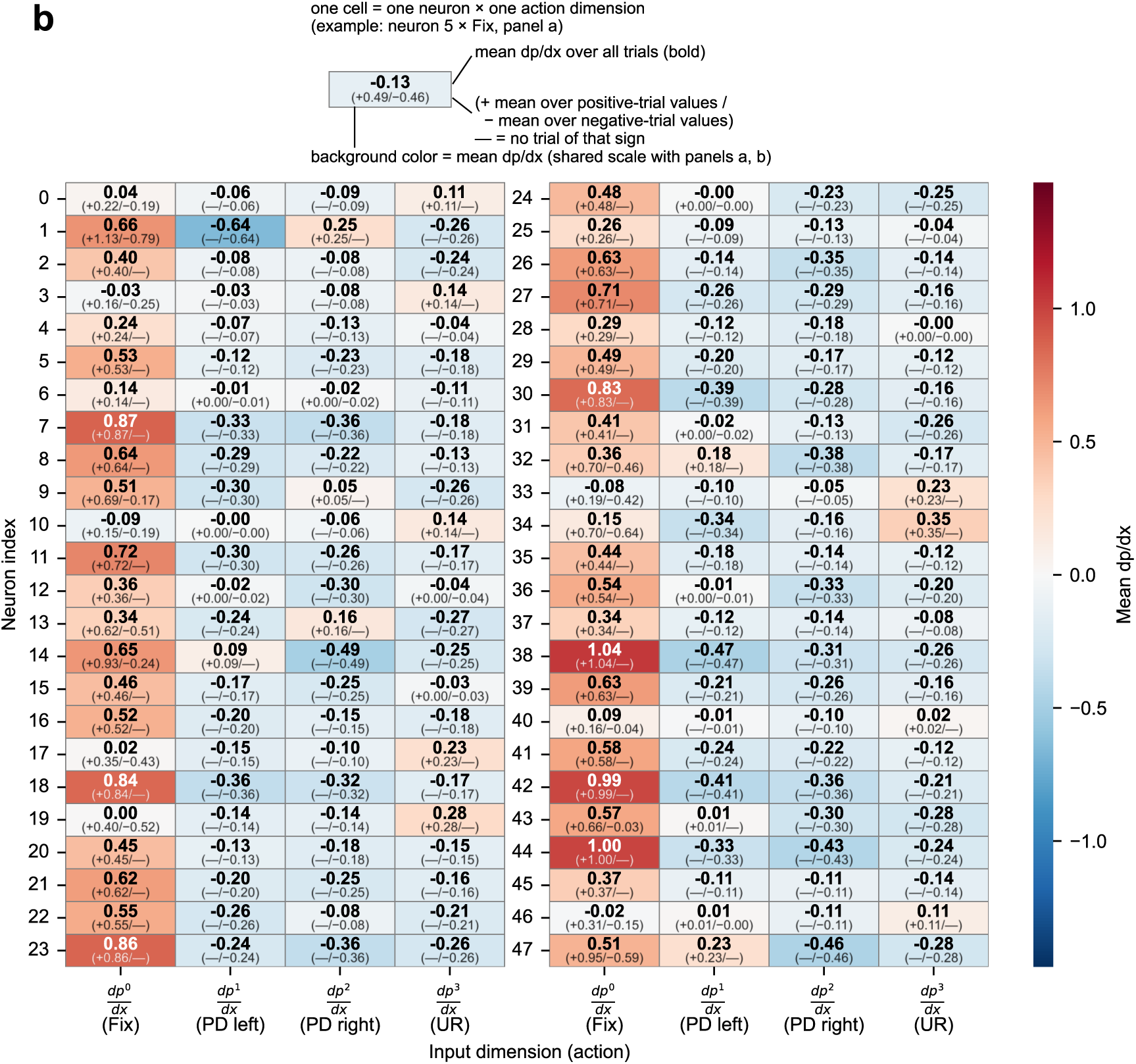
The choice probability contributions for each unit in the SUR agent, computed as described for Extended Data Fig. 11a.

